# Microbial genome functions explain metabolite-driven dysbiosis and *Pseudomonas*-associated ammonium toxicity in *Hydra*

**DOI:** 10.64898/2026.05.18.725911

**Authors:** Natchapon Srinak, Tim Lachnit, Laura Ulrich, Sebastian Fraune, Christoph Kaleta, Jan Taubenheim

## Abstract

Host-associated microbiomes are typically maintained in stable configurations that support host fitness, yet the mechanisms by which metabolic perturbations destabilize these communities remain poorly understood. Using the freshwater cnidarian *Hydra vulgaris* AEP, we systematically assessed microbiome responses to 326 single-metabolite perturbations. Only 17 metabolites, mostly amino acid-related compounds, induced significant compositional shifts in the microbial community. Most shifts are accompanied by transitions from *Curvibacter*- to *Pseudomonas*-dominated or *Legionella*-dominated states, indicating the existence of three alternative community states which can be induced by metabolic triggers. Integrating 16S sequences with functional genomic information, we found that β-diversity strongly predicted functional shifts, whereas reduced α-diversity was associated with loss of metabolic functions. The metabolite perturbations also altered host–microbe interactions, affecting pathogenicity-, glycocalyx-, and nitrogen-related functions. In particular, nitrogen metabolism shifted from ammonia oxidation in *Curvibacter*-dominated communities to ammonia reduction in *Pseudomonas*-dominated states. Experimental validation confirmed that *Pseudomonas* metabolizes L-arginine and drives environmental ammonia accumulation to levels that could impair *Hydra*’s fitness and induce disease phenotypes. Conversely, *Limnobacter* was found to scavenge the environmental ammonia, potentially mitigating the adverse effects. These results demonstrate that metabolite-driven niche reconfiguration can destabilize host-associated microbiomes by coupling compositional shifts to functional change and host pathology, identifying metabolite-driven niche restructuring as a mechanism linking microbial community instability to host dysfunction.

## Introduction

Every eukaryotic organism is colonized by microorganisms that interact closely with the host and contribute to a wide range of physiological processes e.g., immunity, or metabolism, with direct consequences for the evolution of both partners. These integrated relationships are described by the concepts of the metaorganism and the holobiont (1–5). In non-dysbiotic states, these host-microbe interactions typically provide net benefits to the collective in various aspects, for example cell development, nutrient acquisition, and protection against pathogens (6,7). In order to maintain functional microbiomes, the host is able to positively and negatively select its microbes by niche provision and the eradication of potentially harmful microbial members by the immune system (7–11).

However, this homeostasis might be disrupted by environmental perturbations like a change in the trophic environment of the microbiome, thereby changing the ecological niche and masking the host filter (12–15). This can induce dysbiosis, often resulting in detrimental consequences and disease states in the host organism (12,16–20). Consequently, diet is a strong determinant of host-associated microbial composition (21,22). In humans, this translates to lifestyle choices/conditions which determine the microbial community composition. For example, whether a human cohort lives in a non-industrialized or an industrial environment, has a large impact on microbial abundances (23,24), as do dietary choices like high fiber and high fat diets (21,25,26).

Taxonomic alterations are usually associated with changes in the community’s metabolic functional capacity which, in turn, impact host physiology. Consequently, attempts have been made to elucidate general rules governing microbial responses to dietary perturbations and to use controlled dietary interventions as potential strategies to restore microbial homeostasis for the host benefit (27–31). However, the complexity of diet-microbiome, diet-host, microbe-microbe and host-microbe interactions continues to hinder our understanding and demand the usage of suitable and tractable model organisms to trace these interactions.

In this study, we investigated how trophic perturbations affect the host-associated microbiome of *Hydra vulgaris* AEP, which maintains a stable and simple microbiome (32), with only a few key colonizers reported. *Curvibacter*, *Acidovorax*, *Legionella*, *Undibacterium*, *Limnobacter,* and *Pseudomonas* make up the majority of *Hydra*’s microbial community (33–35). *Hydra* uses negative and positive selection to maintain a stable microbiome, by secretion of antimicrobial peptides (AMPs) and the provision of a glycocalyx layer at the epithelial surface (10,11,36–38). In addition, previous studies have demonstrated that the microbiota significantly influences *Hydra*’s physiology e.g., protection against pathogenic infection, regulation of body contractions, and modulation of stem cell differentiation (20,39–42). In summary, *Hydra* shows all hallmarks of a complex metaorganism, while its individual components remain simple enough to trace them experimentally. These characteristics make *Hydra* a relevant model system for dissecting the mechanisms underlying host-microbiome-environment interactions.

Here, we investigated how a large set of single-metabolite perturbations influence the composition and functional responses of *Hydra*-associated microbial communities. We integrated 16S amplicon sequencing with genome-resolved analyses to link community structure and function, improving interpretability of phenotypic experiments. We found that only a small subset of metabolites, predominantly amino acids and their derivatives, induced pronounced shifts in both bacterial diversity and functional profiles, indicating a tight coupling between structure and functional activity. These shifts were largely driven by a transition in community co-dominance from *Curvibacter* to *Pseudomonas* and correlated with pathogenicity, ammonia metabolism and glycocalyx utilization with direct consequences for the host fitness. Finally, we confirmed the functional shift predictions in in vitro and in vivo experiments using the tractable model organism *Hydra*.

## Results

### Amino acid supplementation disturbs *Hydra*-associated microbial composition

We investigated how host-associated microbial composition and function respond to environmental single-metabolite changes, using the freshwater polyp *Hydra vulgaris* AEP as a model organism. We used previously published amplicon sequencing data of *Hydra*-associated microbiomes challenged by 326 single metabolite perturbations utilizing Phenotype MicroArray™ assays (BioLog) representing carbon (PM1 and PM2A), nitrogen (PM3B), as well as phosphorus and sulfur sources (PM4A) (12). In these experiments, *Hydra* was challenged with the medium of the respective plates, thereby exposing the surface-associated microbiome to defined single-metabolite conditions. Metabolite conditions in PM1, PM2A, and PM4A plates were mostly unique whereas some nitrogen conditions in PM3B overlapped with carbon conditions in PM1 and PM2A (Supplementary Figure 1).

Across all 16S amplicon samples, we identified 35 amplicon sequence variants (ASVs) consistently present under different metabolite conditions (Supplementary Table 1, Supplementary Figure 2 and 3). We found a similar microbial composition as previously reported with *Curvibacter* (99.82% prevalence), *Pseudomonas* (93.62% prevalence), *Acidovorax* (99.91% prevalence), *Acinetobacter* (90.11% prevalence), *Legionella* (99.28% prevalence), *Rhodoferax* (99.91% prevalence), and *Limnobacter* (96.76% prevalence) genera, accounted for a median 81.63% relative abundance across samples (Figure 1A, Supplementary Figure 2 and 3). *Curvibacter* was the dominant genus in the control and most experimental conditions. Its relative abundance declined with some metabolite supplementations including amino acids, tricarboxylic acids, fatty acids, and alcohols (Figure 1A and 1B, Supplementary Figure 4 and 5). Among these conditions, *Pseudomonas* frequently emerged as a co-dominant taxon in the microbial community (Figure 1A). Other taxonomic changes included an increase of *Acidovorax* in the presence of flavones and alkylthiols; *Legionella* with fatty alcohols and medium-chain hydroxy acids and derivatives, and *Rhodoferax* under non-metal nitrites and imidazolines (Figure 1A and 1B). We observed that α-diversity varied with the metabolite supplementations, however none of these changes were significantly different from the control conditions (Figure 1C, Supplementary Figure 6 and 7).

**Figure 1:**
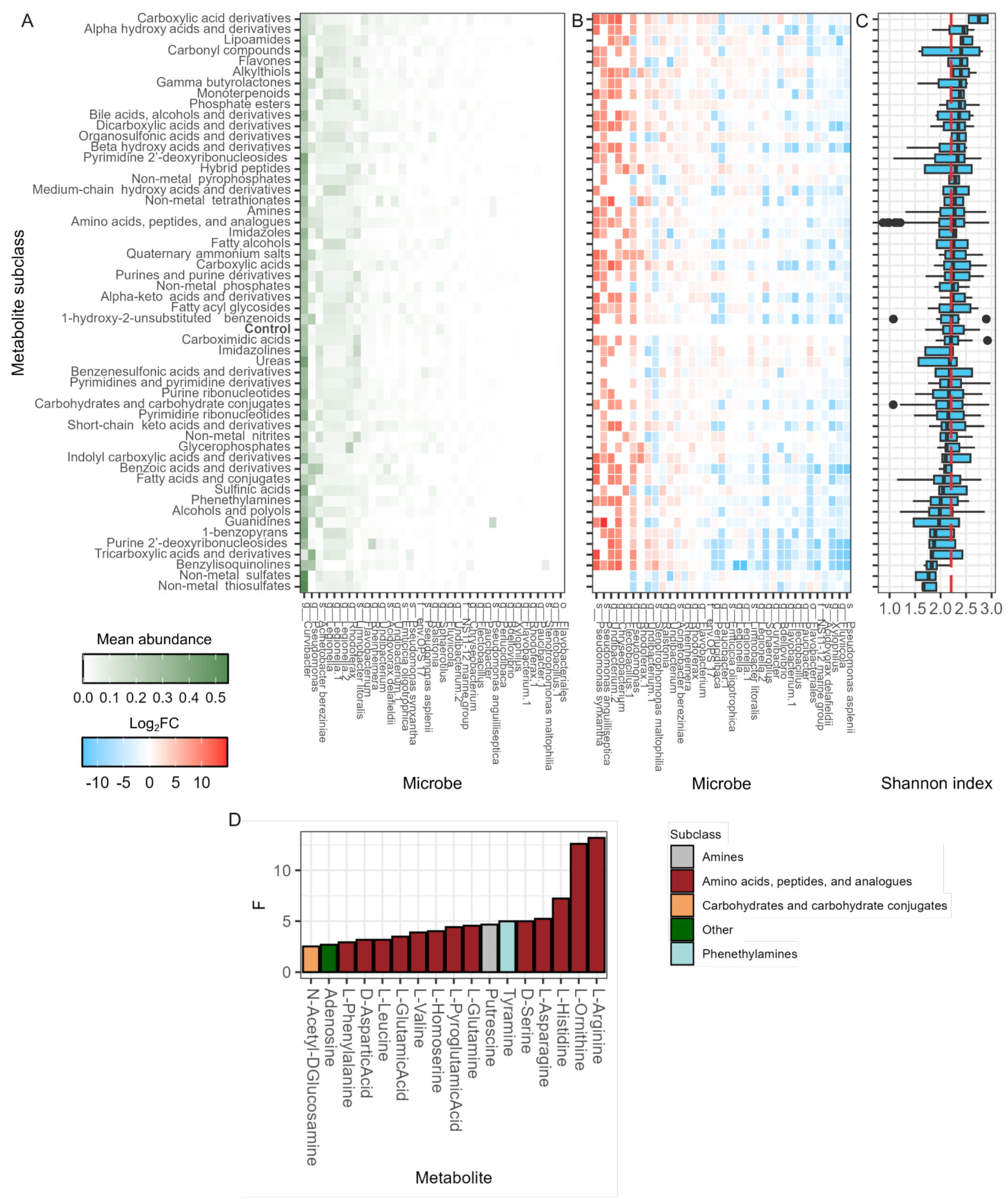
*Hydra*-associated microbiome changes under different metabolite-perturbed conditions. (A) Mean relative abundance of microbes, (B) log 2-fold change of microbial abundance between testing and control conditions, and (C) Shannon α-diversity index in different metabolite subclass conditions (red line indicating median of control conditions). (D) F values from pairwise PERMANOVA indicating the magnitudes of explained variation in Bray-Curtis dissimilarity between metabolite perturbation and the control condition (FDR < 0.05).

To quantify changes in β-diversity due to metabolite supplementation, we calculated the Bray-Curtis dissimilarity index. Testing for changes in the β-dispersion revealed that the variation of microbial composition within metabolite subclasses was significantly different (p = 2.2 × 10⁻¹⁶), whereas no significant differences were observed at the level of individual metabolites (p = 1). For individual metabolites, we performed pairwise PERMANOVA comparisons between each metabolite supplementation and control conditions. Only 17 of 326 (5.21%) of the tested metabolites altered microbial compositions significantly (FDR < 0.05) (Figure 1D), indicating a strong resilience of the microbiome to external perturbation. An enrichment analysis showed that significant compounds were overrepresented in the amino acids, peptides, and analogues subclass (3.09-fold enrichment; hypergeometric test, FDR = 7 × 10⁻⁶). This finding indicates a strong influence of this metabolite group on microbial composition. Further, the amino acids, peptides, and analogues subclass was enriched when testing with metabolites which were significantly associated to Shannon diversity without multiple testing correction (Supplementary Figure 8, hypergeometric test, FDR = 3.34 × 10⁻³).

### *Hydra*-associated microbial species form six functionally different groups

We hypothesized that the changes mediated by different metabolite supplementations were accompanied by changes in the functional capacity of the microbial community. To explore these functional shifts, we assembled 59 metagenomics assembled genomes (MAGs, Supplementary Figure 9 and 10) from a distinct set of genomic samples and 6 genomes from bacterial isolates (see Methods for details). To combine the genomes with our perturbation experiment, we developed a Method for linking 16S rDNA sequences directly to reference genomes (Supplementary Figure 11). In the first step, we mapped the metagenomic sequences to reconstructed genomes and 16S ASVs and used the information of commonly mapped reads to link genomes to ASVs. In a second step, we predicted 16S sequences from the metagenomic reads using specific 16S assembly (MATAM (43)), from the genomic sequences themselves (barrnap https://github.com/tseemann/barrnap) or from the taxonomic associations to the genome (GTDBTk (44,45)). Afterward, we aligned these predicted 16S sequences to the ASVs from the amplicon sequencing and linked both with a 95% similarity cutoff (Supplementary Figure 12). With this, we were able to successfully link 27/35 ASV to 17 individual genomes (Supplementary Table 3) enabling a deeper investigation of abundance-weighted community-level functional activity after metabolite perturbation. The redundant linkage indicates closely related clades within the *Hydra*-associated microbiome.

The successfully linked ASVs accounted for medians of 92.5%, 92%, 82.5%, and 85.3% relative abundance of the microbial community in PM1, PM2A, PM3B, and PM4A, respectively (Figure 2A).

**Figure 2:**
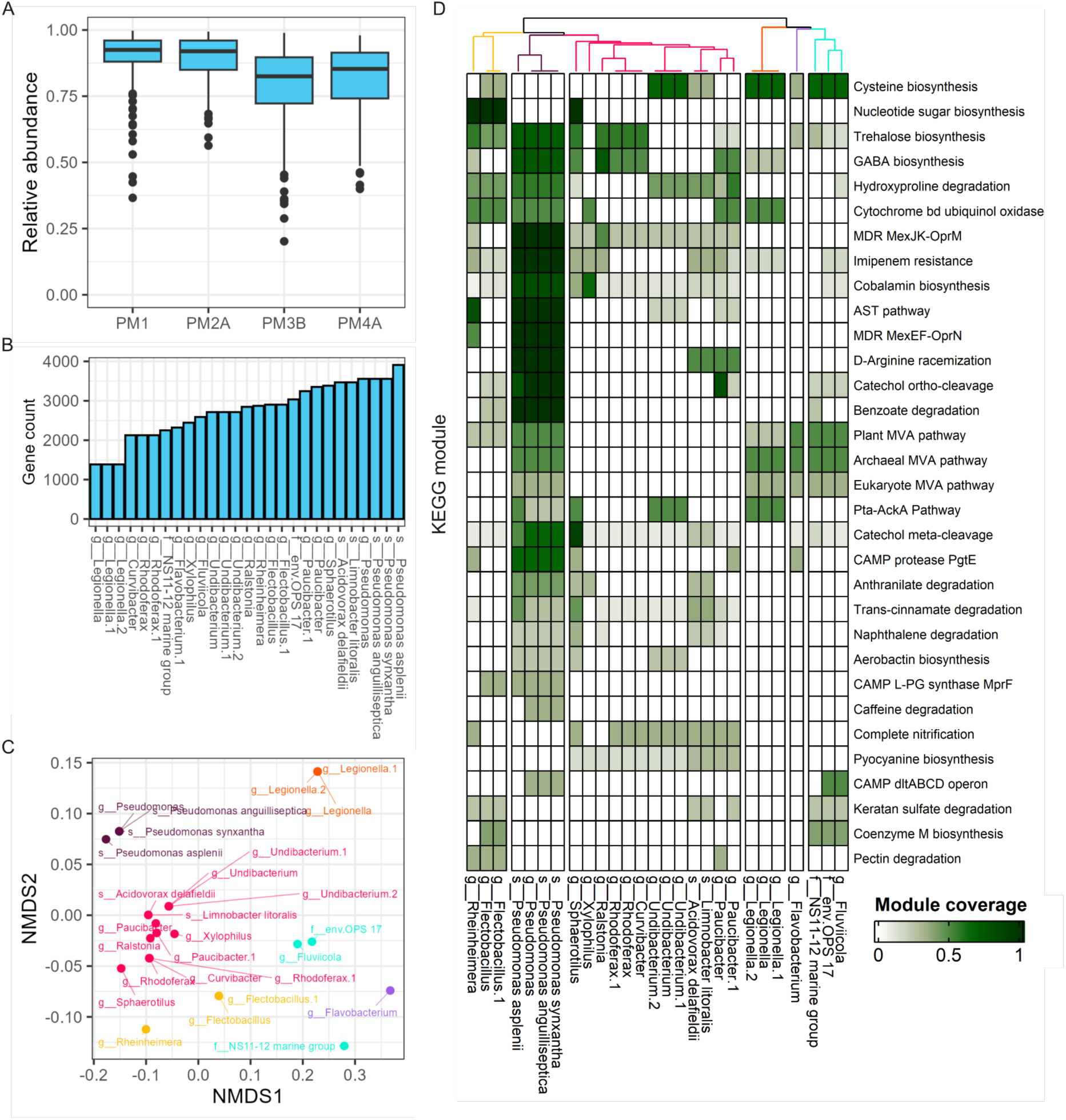
Functional module coverage of individual microbes. (A) Relative abundances of microbial taxa after 16S–genome linkage. (B) Total number of KEGG metabolic and signalling genes assigned to each microbe. (C) NMDS ordination of microbial functional coverage profiles, where colors indicate groups of functional similarity. (D) Heatmap of selected functional modules highlighting coverage patterns across microbes. Modules displayed were identified as significantly different among groups (Kruskal-Wallis test, adjusted p < 0.05) and further filtered by functional variability criteria (Gini coefficient > 0.5 and standard deviation > 0.1). AST, arginine succinyltransferase; CAMP, cationic antimicrobial peptide; L-PG, lysyl-phosphatidylglycerol; MDR, multidrug resistance; MVA, mevalonate; Pta-AckA, phosphate acetyltransferase-acetate kinase.

The individual microbial genomes were annotated for their functional potential, and the metabolically related genes were used to quantify the functional coverage of each KEGG module (see Methods for details, Supplementary Table 4 and 5). Firstly, we observed that the number of metabolic and signalling genes varied across *Hydra*-associated microbial species (Figure 2B). Genomes of microbes such as *Pseudomonas* spp., *Sphaerotilus*, *Limnobacter*, and *Acidovorax* showed higher numbers compared to that of other microbes. In contrast, *Flavobacterium, Fluviicola* and *Undibacterium* showed intermediate gene numbers, while *Curvibacter*, *Rhodoferax*, and *Legionella* had the lowest. Using the module coverage profiles, we grouped the microbial species into six distinct functional clusters (Figure 2C). The largest cluster included several dominant colonizers of *Hydra*, such as *Curvibacter*, *Rhodoferax*, *Acidovorax*, and *Undibacterium*, indicating shared functional profiles and possible ecological niche overlap (*Curvibacter* group, Figure 2C, pink). In addition, *Pseudomonas* (*Pseudomonas* group, Figure 2C, plum), *Legionella* (*Legionella* group, Figure 2C, orange), and *Flavobacterium* (*Flavobacterium* group, Figure 2C, violet) formed individual genus clusters. *Flectobacillus* and *Rheinheimera* grouped together (*Reinheimera* group, Figure 2C, yellow), as did *Fluviicola*, NS11-12, and env.OPS17 (*Fluviicola* group, Figure 2C, cyan). On the phylogenetic level, the 16S sequences of *Curvibacter*, *Rhodoferax*, and *Acidovorax*, showed high similarity, while *Undibacterium* showed higher divergence, yet was more similar to the three other genera in this cluster than any other ASV found (Supplementary Figure 12). Similarly, *Fluviicola*, NS11-12 and env.OPS17 had related 16S sequences in our analysis. In contrast, 16S similarity was very low between *Flectobacillus* and *Rheinheimera*, indicating that functional similarity does not necessarily reflect phylogenetic similarity in this group. In general however, our functional clustering is supported by phylogenetic signals in the 16S gene (Supplementary Figure 12, Supplementary Table 3).

These patterns suggest that functional redundancy within clusters could buffer the microbiome against fluctuations in the abundance of individual species, thereby helping to preserve overall community function. In contrast, shifts in abundance across distinct functional clusters may be more likely to alter the functional output of the entire community, potentially resulting in a change in community structure.

We next investigated the specific functional modules that distinguish the microbial species associated with *Hydra* (Figure 2D). In the *Curvibacter* group, several unique modules with higher coverage were identified, including complete nitrification, and pyocyanin biosynthesis, while isoprenoid biosynthesis modules (mevalonate pathway) were missing. The *Pseudomonas* group harbored multiple antibiotic resistance modules, such as multidrug resistance, imipenem resistance, and cationic antimicrobial peptide (CAMP) resistance. Additionally, genes involved in aerobactin biosynthesis, a siderophore facilitating iron scavenging from the environment and potentially the host (46,47), were more prevalent, indicating a pathogen-like lifestyle. This aligns with previous findings indicating that *Pseudomonas* competes for iron with the host, which could mediate pathogenicity and increase host mortality (12). Metabolically, *Pseudomonas* also exhibited a specialization in arginine metabolism, but a relative deficiency in cysteine biosynthesis compared to the *Fluviicola*, *Legionella*, *Rheinheimera*, and *Flavobacterium* groups. Interestingly, genes related to keratan sulfate degradation were more abundant in the *Fluviicola* and *Rheinheimera* groups and some taxa in the *Curvibacter* group. This function is associated with the breakdown of glycosaminoglycans, a component of the *Hydra* glycocalyx (48). Our findings suggest that these groups may act as primary colonizers and degraders in the *Hydra* microbiome, extracting carbon sources from host-derived glycoconjugates and potentially supporting the broader microbial community through shared metabolic products.

### Taxonomic diversity partially reflects functional diversity

We further investigated changes of microbial community functions after single-metabolite perturbations. We quantified the abundance of each functional module in response to metabolite supplementation and compared them to the control conditions (see Methods for details, Supplementary Table 6 and 7). Among all 326 tested metabolites, only 38 showed at least one significant functional module abundance change, confirming the stability of the microbiome against metabolic perturbations also on the functional level. The metabolites that induced most significant functional changes belonged to the class of alcohols and polyols; fatty acids and conjugates; benzoic acids and derivatives; amino acids, peptides and analogues, or amines (Figure 3A). In turn, most of the functional change inducing metabolites fell in the subclass of amino acids, peptides and analogues, followed by fatty acids and conjugates, and carbohydrates and carbohydrate conjugates (Figure 3B). As expected, taxonomic change in the microbiome structure was directly associated with functional change in the community. Notably, Bray-Curtis distance from the control condition explained 24.7% of the variance in the number of significantly altered functional modules (FDR = 4.14 × 10⁻²¹) (Figure 3C). Furthermore, we calculated the absolute difference in α-diversity metrics to control conditions and found that evenness and Shannon diversity significantly explained 7.2% (FDR = 1.66 × 10⁻⁶) and 3.4% (FDR = 1.10 × 10⁻³) of the variance in the number of changed functional modules respectively, whereas richness was not significantly associated (Figure 3C). These findings suggest that both diversity measures can serve as a rough proxy for functional changes in the microbial community, though they represent only a fraction of the functional variability. A higher number of altered modules was linked to lower microbial diversity, and especially increased the numbers of down-regulated functions (Figure 3D-E). This suggests that α-diversity is directly associated with functional diversity in the community. Several metabolite conditions, such as itaconic acid, hydroxy-L-proline, quinic acid, hydroxylamine, and D-asparagine, were associated with the highest number of functional changes. These conditions also exhibited a concurrent decline in Shannon-diversity, supporting the hypothesis that reductions in α-diversity may reflect the loss in microbial community function with a reduction of host-associated microbiome diversity.

**Figure 3:**
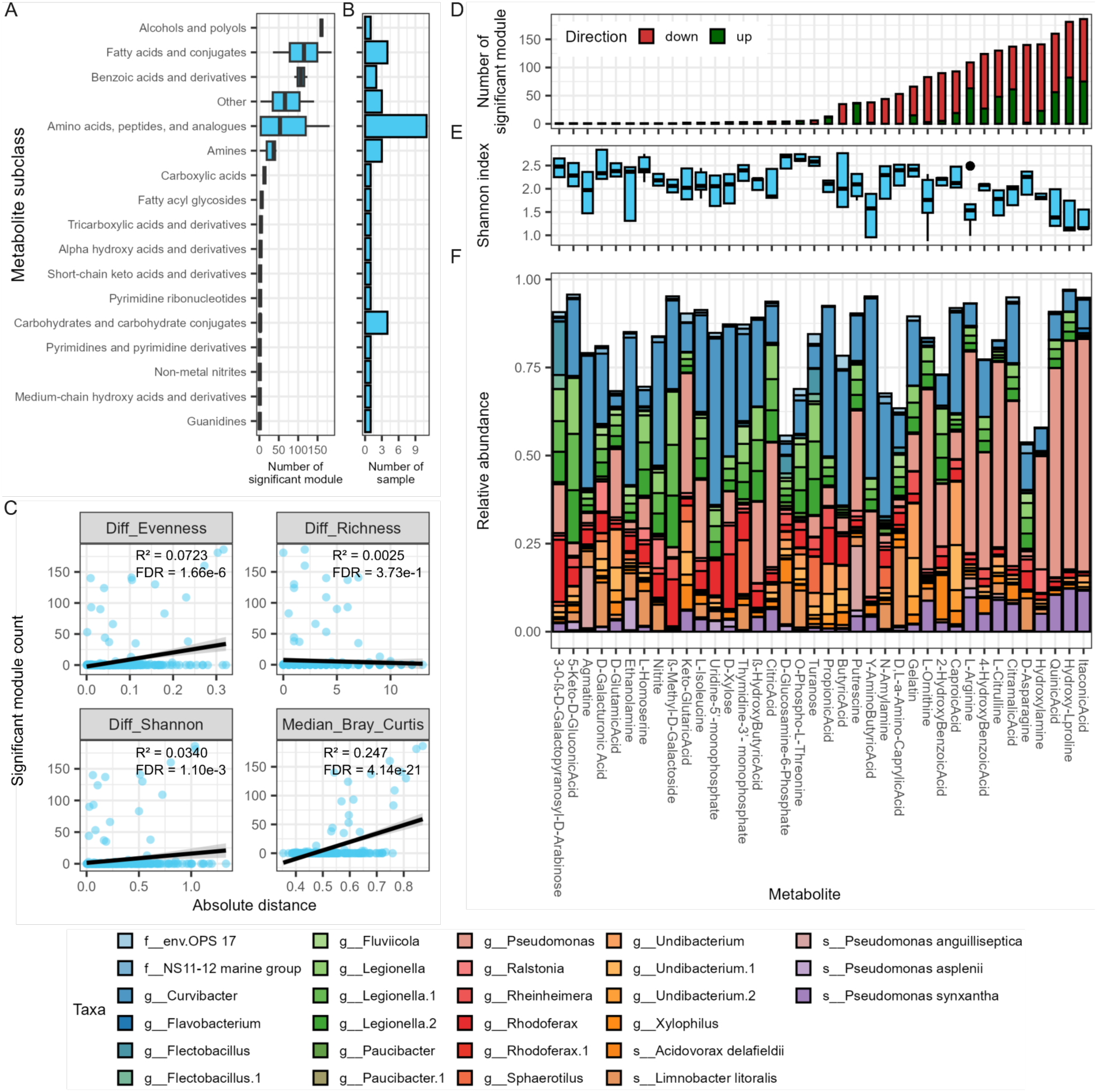
Microbial functional changes associated with community structural alterations. (A) Number of significantly changed functional modules across metabolite subclasses. (B) Number of samples containing at least one significantly changed module for each metabolite subclass. (C) Associations between changes in functional modules and microbial community structure indices (evenness, richness, Shannon diversity, and Bray-Curtis dissimilarity). Summary of significant changes in (D) functional modules, (E) Shannon diversity, and (F) microbial profiles across metabolite conditions exhibiting at least one significantly changed module.

### *Pseudomonas* dominance restructures microbiome function

The *Curvibacter* group dominated in abundance (*Curvibacter* dominance) in 267 conditions (including the control condition), followed by the *Legionella* group (*Legionella* dominance) in 35 conditions and the *Pseudomonas* group (*Pseudomonas* dominance) in 24 conditions. These dominance-states represented three exclusive states in the microbiome composition (Supplementary Figure 13), which are induced by specific metabolite perturbations. Although observed for several metabolite conditions (including xylitol, β-methyl-D-xyloside, stachyose, N-acetyl-neuraminic acid, and D-fucose), *Legionella* dominance caused relatively little perturbation to the overall microbial community function (Figure 3D-F, Supplementary Figure 13, boxplot). On the other hand, *Pseudomonas* dominance exhibited the largest number of significantly affected modules per condition (Figure 3D-F). Furthermore, the relative abundance of the *Pseudomonas* group is a good predictor for a functional change in the community (logistic regression, p = 0.0154), while the abundances of *Curvibacter* and *Legionella* showed no significant associations (Supplementary Figure 13, dotplot). In addition to *Pseudomonas*-associated functional changes, we observed larger numbers of significantly altered modules when supplementing D-asparagine, caproic acid, gelatin, D-L-ɑ-amino caprylic acid, and N-amylamine. These changes occurred alongside shifts in less abundant species, such as *Undibacterium*, *Acidovorax delafieldii*, and *Limnobacter litoralis* from the *Curvibacter* group (Figure 3D-F). In general, communities losing many functional modules typically showed reduced relative abundances of taxa such as *Curvibacter*, *Fluviicola*, *Legionella*, and *Rhodoferax* (Figure 3F). This finding suggests that these microbes may contribute to a majority of unique functions in the *Hydra* microbiome, which appear to diminish under stress or invasion. Such disruptions may lead to overgrowth of opportunists, community-level dysbiosis, or even host mortality as demonstrated previously (12).

Among the metabolites whose supplementation elicited significant functional differences, several were metabolically related compounds and were associated with corresponding shifts in microbial composition (Figure 3F). For instance, L-citrulline can act as a precursor to L-arginine, while L-ornithine, produced via hydrolysis of L-arginine, can undergo decarboxylation to form putrescine (Supplementary Figure 14). Similarly, nitrogenous compounds such as hydroxylamine can be reduced to ammonia, which serves as a nitrogen source for the biosynthesis of several amino acids, including L-arginine. Moreover, L-ornithine can be metabolized into L-glutamate-5-semialdehyde, which can subsequently be converted into L-proline and hydroxy-L-proline (Supplementary Figure 14). These metabolically connected amino acid conditions were consistently associated with an enrichment of *Pseudomonas*, replacing *Curvibacter* as the dominant colonizer under control conditions. In contrast, gelatin supplementation supported *Undibacterium*, despite gelatin being rich in L-proline and hydroxy-L-proline, compounds that favor *Pseudomonas* growth. This suggests that additional factors, such as the presence of other compounds in gelatin, differential substrate uptake preferences, or varying degradation capacities among microbes and host, may influence microbial community structure, underscoring the complexity of microbial adaptation in host-associated environments. Additionally, metabolite conditions involving butyric acid, propionic acid, and caproic acid (all fatty acids) tended to enrich *Curvibacter*, *Undibacterium*, and *Rheinheimera*, but not *Pseudomonas*. In contrast, itaconic acid and citramalic acid, relating to intermediates of the tricarboxylic acid (TCA) cycle, were associated with increased *Pseudomonas* group abundance. These observations suggest that even within the same chemical category, such as organic acids, structural and metabolic differences can lead to distinct microbial community compositions, while metabolically closely related compounds tend to induce more similar shifts in microbial communities. Accordingly, when such related metabolites shape similar microbial profiles, the resulting functional responses are also likely to change in a coordinated manner.

### Functional responses are directly linked to metabolite perturbations

We observed that functional changes could be broadly categorized into two groups, primarily depending on the dominant microbial group: *Curvibacter* or *Pseudomonas* (Figure 4A, Supplementary Figure 15 and 16). In samples with *Pseudomonas* dominance, most significantly changed modules were reduced in abundance, indicating the reduction of functional diversity. On the other hand, we, observed significant increases in the functional abundance of several modules, such as D-arginine racemization, the arginine succinyltransferase pathway, hydroxyproline degradation, ornithine biosynthesis, ornithine-ammonia cycle, and GABA biosynthesis, in response to the addition of L-arginine and closely related compounds such as L-citrulline, putrescine, hydroxylamine, and hydroxyproline (Figure 4A and Supplementary File 1). These modules are either directly involved in, or closely connected to the metabolic pathways of the supplemented compounds (Supplementary Figure 14), promoting bacterial species harboring these functions. For example, the *Pseudomonas* group genomes encode for D-arginine racemization and the arginine succinyltransferase (Figure 2D) and showed a significant abundance increase with L-arginine supplementation, linking environmental conditions, functional capacity and bacterial abundance in a host associated context (Figure 4A and Supplementary File 1). Furthermore, supplementing certain organic acids, such as TCA cycle intermediates (e.g., itaconic acid and citramalic acid) and a cyclitol (quinic acid), also triggered functional responses similar to supplementation with L-arginine related compounds, inducing the same *Pseudomonas* dominance. However, these metabolites cannot be directly linked mechanistically to induced metabolic functions as seen in the L-arginine-related supplementations, however, they may serve as metabolites for L-arginine-related pathways. In contrast, propionic acid tended to promote the *Curvibacter* group accompanied by the increase in several amino acid biosynthesis modules, like proline biosynthesis, urea cycle, lysine biosynthesis, and tyrosine biosynthesis (Figure 4A and Supplementary File 1).

**Figure 4:**
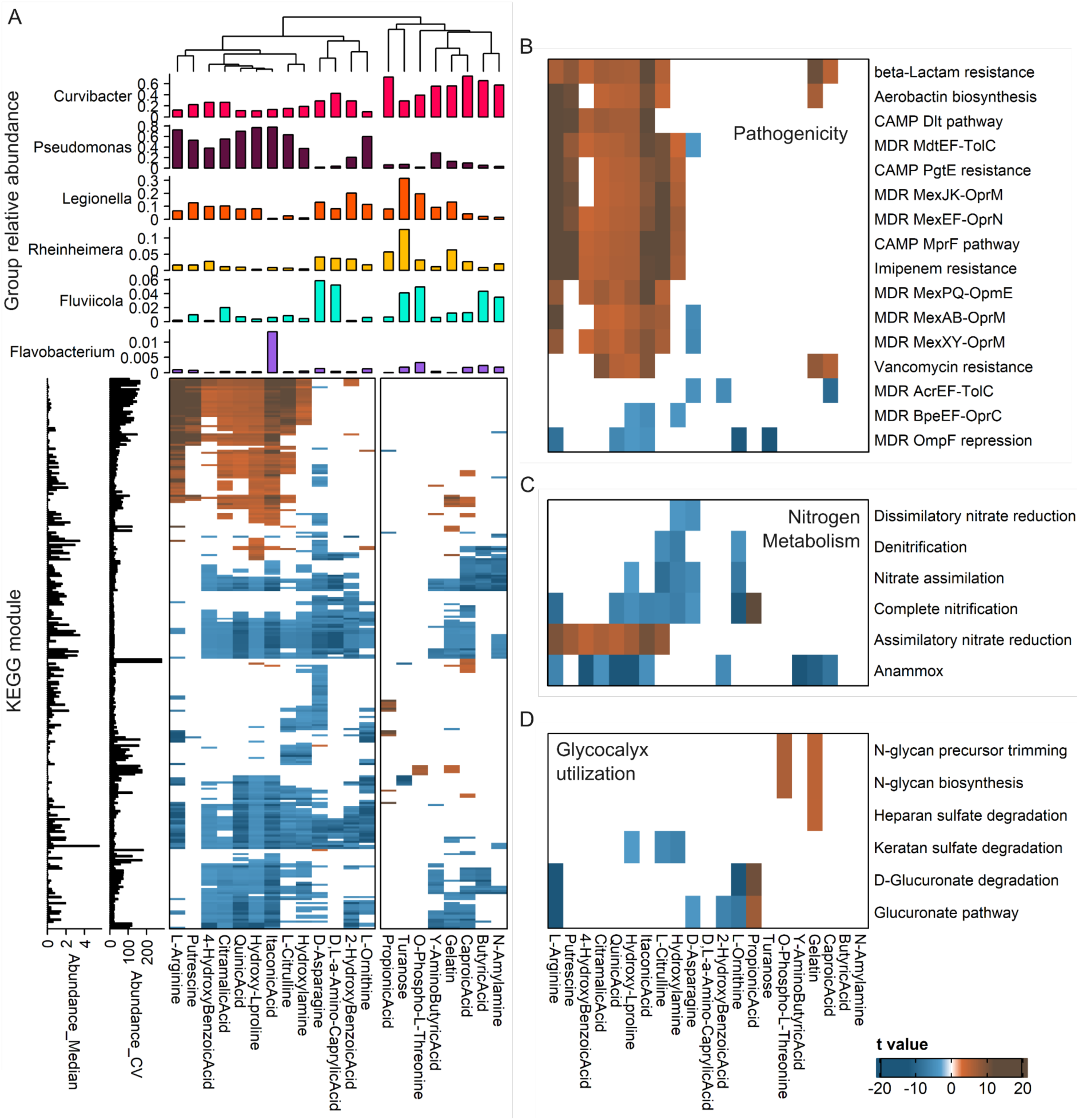
Heatmap of community functional abundance changes under different metabolic perturbations. (A) Overall profiles of differential community KEGG module abundance across all tested conditions (FDR < 0.05). The heatmap clusters into two major groups, largely corresponding to patterns of microbial group relative abundances (top). Median and coefficients of variation (CV) for each module abundance are indicated alongside the rows. Panels (B-D) show enlarged views focusing on three functional systems potentially related to host-microbe interaction: (B) pathogenicity, (C) nitrogen metabolism, and (D) glycocalyx utilization. CAMP, cationic antimicrobial peptide; Dlt, D-alanylation of lipoteichoic acids; MDR, multidrug resistance.

### Metabolite perturbations affect *Hydra*-microbe interactions

L-arginine and related metabolites, TCA cycle intermediates, and cyclitols promoted functions linked to microbial competitiveness and pathogenic potential. These included multidrug resistance efflux systems (e.g., MdtEF-TolC, MexEF-OprN, MexJK-OprM, and MexAB-OprM), as well as resistance mechanisms to imipenem, cationic antimicrobial peptides (CAMPs), and β-lactams (Figure 4B). The increased abundance of these functions coincided closely with an elevated relative abundance of the *Pseudomonas* group (compare Figure 2D and 4B). Moreover, the aerobactin biosynthesis module, known to enhance microbial competitiveness, was enriched under these conditions, posing a potential risk of increased pathogenicity to the host. Functional modules related to *Hydra* glycocalyx degradation, including keratan sulfate degradation, the uronate pathway, and D-glucuronate degradation, were consistently reduced under L-arginine and TCA cycle intermediate conditions (Figure 4D). Keratan sulfate degradation was linked to the *Rheinheimera*, *Fluvicola* and *Curvibacter* group (Figure 2D) and glycocalyx degradation pathways were strongly increased under propionic acid, O-phospho-L-threonine, and gelatin conditions, where these bacterial groups showed increased relative abundances (Figure 4A, Supplementary Figure 15 and 16).

Furthermore, we observed a decrease in ammonia oxidation pathways (dissimilatory nitrate reduction, denitrification, complete nitrification, anammox) and an increase in the assimilatory nitrate reduction module in response to L-arginine, TCA intermediates, or caproic acid supplementation (Figure 4C). This indicates a shift from ammonia degradation to ammonia production under these conditions. Specifically, complete nitrification is linked with the genome function of the *Curvibacter* group (Figure 2D) while assimilatory nitrate reduction is associated with increased abundances of the *Pseudomonas* group (Figure 4A and 4C, Supplementary Figure 13, barplot). Hence, we argue that under normal conditions, the *Curvibacter* group utilizes ammonia, potentially cross-fed by the *Hydra* host, as an electron donor. Following L-arginine supplementation, *Pseudomonas* dominance shifted ammonia metabolism toward excessive microbial secretion as nitrogenous waste. This transition might reduce metabolic host-microbe dependency, leading to ammonia accumulation that could be toxic to *Hydra* and contribute to previously reported Pseudomonas pathogenicity (12).

### *Pseudomonas* group shows increased ammonia release

To experimentally confirm our analysis linking ammonium metabolism to metabolite perturbation and community shifts, we examined the ability of *Pseudomonas* to generate ammonia under the L-arginine condition, and of *Curvibacter* group members to oxidize ammonia. We measured ammonium levels in in vitro cultures after supplementation with either NH_3_ or L-arginine in an otherwise nutrient free medium (S-Medium). Further, we tested the microbial production or consumption of NH_3_ in mono- or dualcolonized *Hydra* in S-Medium and in supplementations with ammonia or L-arginine (Figure 5).

**Figure 5:**
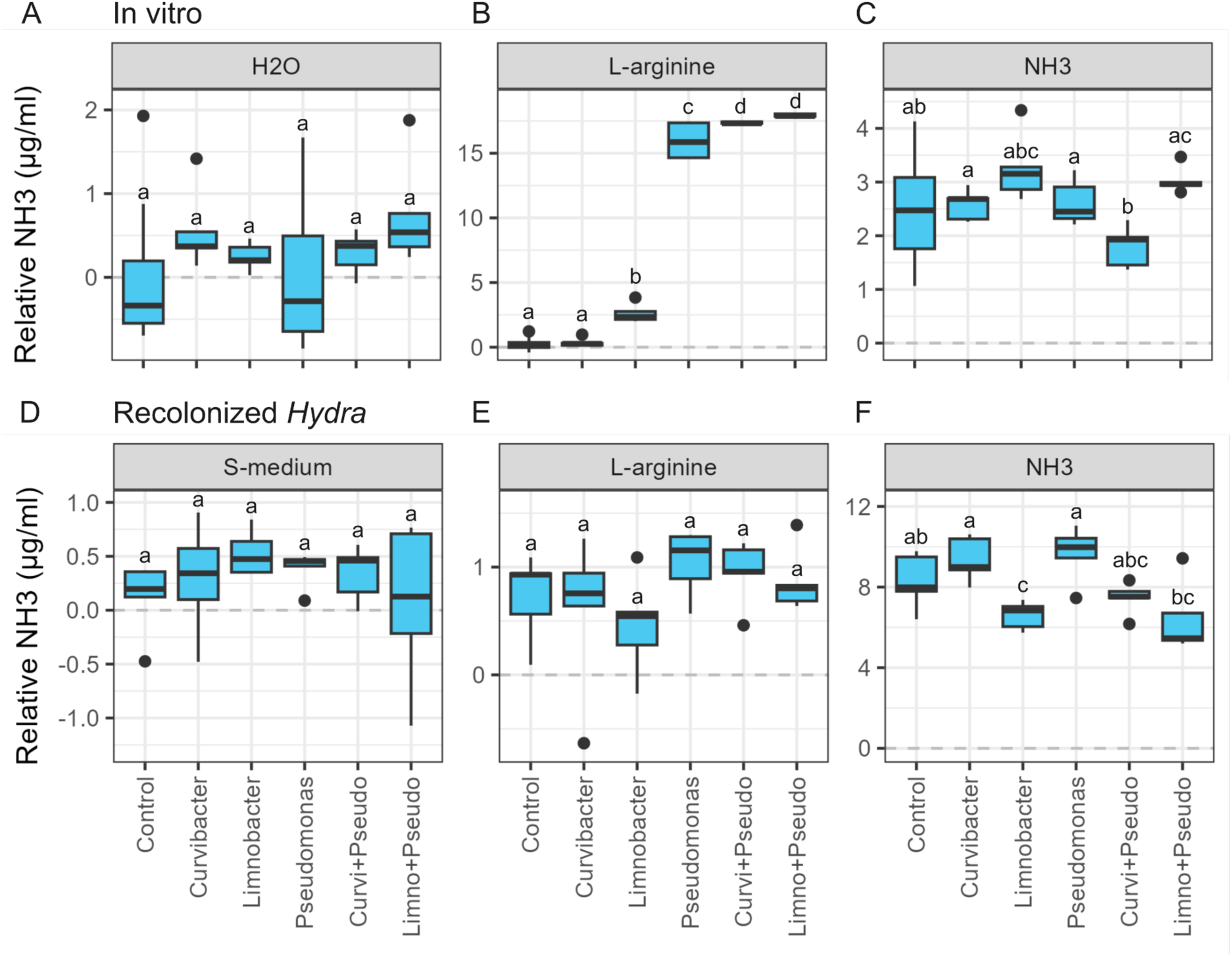
Ammonia uptake and secretion by microbes in *Curvibacter* and *Pseudomonas* groups. Ammonia concentrations measured in *in vitro* experiments in water only (A), or after addition of L-arginine (B), or NH₃ (C). Ammonia concentrations measured in recolonized *Hydra* cultured in S-medium (D), L-arginine (E), and NH₃ (F) conditions. Letters above boxplots indicate statistically significant different groups (t-test, FDR < 0.05).

In vitro experiments demonstrated that *Pseudomonas* can metabolize L-arginine and release ammonia as a metabolic by-product in mono- or dual-culture together with *Curvibacter* or *Limnobacter* (also *Curvibacter* group, t-test, FDR < 0.05, Figure 5B, Supplementary Table 8). Supplemented ammonia concentrations remained unchanged to control conditions in the *Pseudomonas* culture, indicating no ability of *Pseudomonas* to reduce ammonia in vitro (Figure 5C). However, the dual-culture of *Pseudomonas* and *Curvibacter* significantly reduced the ammonia concentrations compared to both mono-culture conditions, indicating an interaction between both bacteria leading to a slight, but net reduction of ammonium (t-test, FDR < 0.05, Figure 5C). L-arginine supplementation did not induce any significant change in ammonia concentrations in vivo (Figure 5E). This is in stark contrast to *Pseudomonas* in vitro conditions and suggests a buffering effect of the *Hydra* host on the NH_3_ levels with *Pseudomonas* colonization. However, we noted a non-significant trend of increased ammonia levels in *Pseudomonas* mono-colonization and a reduction in ammonia concentrations when *Limnobacter* was mono- or dual-colonized (Figure 5E). The latter trend was confirmed when supplementing in vivo cultures with ammonium, here we observed reduced levels of ammonium in *Limnobacter* mono- and *Pseudomonas-Limnobacter* dual-colonizations (Figure 5F). *Limnobacter*-*Hydra* interactions seem to induce the nitrification response in *Limnobacter*, which was not observed in vitro.

As hypothesized, we observed that *Hydra* released ammonia into the environment in nutrient-free conditions (Supplementary Figure 17). This might serve as a nitrogen and energy source for microbes in the *Curvibacter* group (e.g., *Limnobacter*) and facilitates colonization under unperturbed conditions. Exposure to high ammonia is toxic to *Hydra* (Supplementary Figure 18), which suggests that the observed buffering after L-arginine supplementation inflicts a fitness burden on the host (and has been observed elsewhere [Tim’s paper]). In vitro ammonia production by *Pseudomonas* under L-arginine supplementation easily reached toxic NH_3_ concentrations for *Hydra* (compare Figure 5B and Supplementary Figure 18), indicating a detrimental threat of *Pseudomonas* to *Hydra* on a metabolic level.

## Discussion

The *Hydra*-associated microbiome exhibited alterations in both community composition and function following single-metabolite perturbations. Overall, we found relatively few metabolites that disturbed the *Curvibacter* dominated microbial community composition (59/326, 18.1%) supporting the previous notion of a stable *Hydra*-associated microbiome under laboratory conditions (34,36). In natural *Hydra* populations, microbiota composition varied substantially and was dominantly shaped by the habitat (49), whereas additional environmental factors such as water temperature and additional colonization from environmental microbes likely influenced microbial community composition (50,51). Our findings are consistent with this ecological observation, suggesting that the microbiome is resilient to many compounds in the absence of additional biotic and abiotic factors, however, defined nutrients and metabolites can function as selective drivers, able to reshape community composition. In this study, we showed that metabolites belonging to the amino acids, peptides, and analogues subclass frequently led to substantial compositional and functional alterations (Figure 1E and 3A). Although from the same category, the effects of these metabolites to microbial characteristics were diverse: for instance, O-phospho-L-threonine increased α-diversity, whereas hydroxy-L-proline reduced it (Figure 1D). D-aspartic acid and glycyl-L-methionine tended to promote the *Curvibacter* and *Legionella* group, while L-arginine and hydroxy-L-proline induced the predominance of the *Pseudomonas* group (Figure 3F, Supplementary Figure 16). In contrast, we found that microbiome responses appeared to depend more on metabolic similarity than on chemical classification. Closely related compounds such as L-arginine, L-ornithine, hydroxy-L-proline (amino acids, peptides, and analogues), and putrescine (amines) produced comparable community structures dominated by *Pseudomonas*, as did TCA intermediates like itaconic acid (fatty acids and conjugates) and citramalic acid (organic acids and derivatives) (Supplementary Figure 14, Figure 3F). These patterns suggest that members of the community exhibit metabolic specialization, responding preferentially to a set of related metabolic precursors.

Across several perturbation conditions, we observed a consistent shift in relative dominance from the *Curvibacter* group to the *Pseudomonas* group. Both are consistently found in the *Hydra* host (20,35). Under normal lab conditions, *Curvibacter* is typically the main colonizer and demonstrated a strong impact on host fitness (20,52), whereas *Pseudomonas* remains at low abundances (35). Both species exhibit a high degree of metabolic niche overlap (53), yet *Curvibacter* is able to outcompete *Pseudomonas* under standard conditions. Though *Curvibacter* is the main colonizer in *Hydra*, it can be outcompeted by *Duganella* (also *Curvibacter* group with similar niche overlap) in dual-colonization experiments (54), implying that other microbe-microbe and host-microbe interactions are needed to stabilize the *Curvibacter* dominance. Our results show that environmental challenges can destabilize these interactions inducing a dramatic change in the microbial community composition, leading to *Pseudomonas* outgrowth.

Further, we could link metabolic pathways associated with these changes. The complete nitrification module was enriched in several microbes in the *Curvibacter* group (Figure 2D and 4D) which serve as an important way to obtain energy from ammonium, a potential host waste product (2) (Figure 6). Nitrification has been previously described as an energy source for microbes and is an important process for host-microbe interactions (55–58). In addition, the *Curvibacter* group, as well as the *Reinheimera* and *Fluvicola* groups, possessed the ability to utilize glycocalyx-related compounds (e.g., N-glycan, heparan sulfate, keratan sulfate, and glucuronate) (Figure 2D and 4D), indicating a certain degree of host-adaptation and -selection as the host potentially produces these specific glycans (59,60). The outermost layer of *Hydra’s* glycocalyx is colonized by microbes i.e., *Curvibacter* (20), which may allow for potential nutrient exchanges between host and microbe. We suspect that these bacterial groups serve as primary degraders of the complex carbohydrates in the glycocalyx, thereby generating a metabolic niche for the other bacteria in the community. This is consistent with previous observations showing that *Curvibacter* colonizes early during the first 1-2 weeks after hatching (35). The metabolites promoting *Pseudomonas* growth could serve as alternative electron donors, and carbon and nitrogen sources (61), which might render some of the existing filters created by the *Curvibacter* interactions with the host ineffective. L-arginine has been described to induce pathogenic phenotypes in *Pseudomonas* (61–63), reflected in the increase of virulence-associated functions, such as multidrug/antibiotic resistance and siderophore genes in our analysis (Figure 4B). Among the pathogen-associated genes, aerobactin production has previously been suggested to impair *Hydra* host health (12), and likely disrupt the host selection filters further. Alternatively, L-arginine has been described to induce a sessile lifestyle and biofilm formation in *Pseudomonas aeruginosa* (63,64). This resembles a similar mechanism in *Curvibacter* where quorum quenching of *Hydra* induced a sessile lifestyle, promoting *Curvibacter* colonization (65). Metabolically, the microbial community shifted functionally, favoring the reduction of nitrogen compounds to ammonia over ammonia oxidation for energy (Figure 4C). This metabolic switch suggests that the community is now supplemented with external energy sources and no longer relies on host-derived compounds for energetic purposes. Instead, the host must detoxify excess ammonium, increasing metabolic demand and thereby reducing host fitness (Figure 6). Freshwater invertebrates often couple their osmoregulation to the excretion of waste ammonia (66), which is hindered with increased environmental NH_3_-levels. In addition, ammonia showed direct toxic effects on the neuroendocrine regulation of several invertebrates (67), which is central to the production and release of AMPs in *Hydra* for microbiome control (11,37,68). Thus, we believe that the observed changes arise from a combination of impaired host selection and metabolite-induced changes in microbial metabolic interactions. In our and previous assays (12) we cannot distinguish between the effect of virulence factor expression and NH_3_ production of *Pseudomonas* causing the health decline in *Hydra* (both seem plausible). Hence we argue that the NH_3_-metabolism of *Pseudomonas* is part of its virulence and weakens the host’s defenses. In other systems, like the coral holobionts, the disruption of nitrogen exchange between host, symbiotic algae, and associated microbes drive bleaching events (69). Ammonia pollution is also known to induce microbial shifts in diverse aquatic animals (70), positioning ammonium homeostasis as an important health factor in aquatic host-microbe interactions. Consistent with this, our results support the view that nitrogen flux constitutes a key axis of host-microbe interactions that require tight regulation to maintain functional stability.

**Figure 6:**
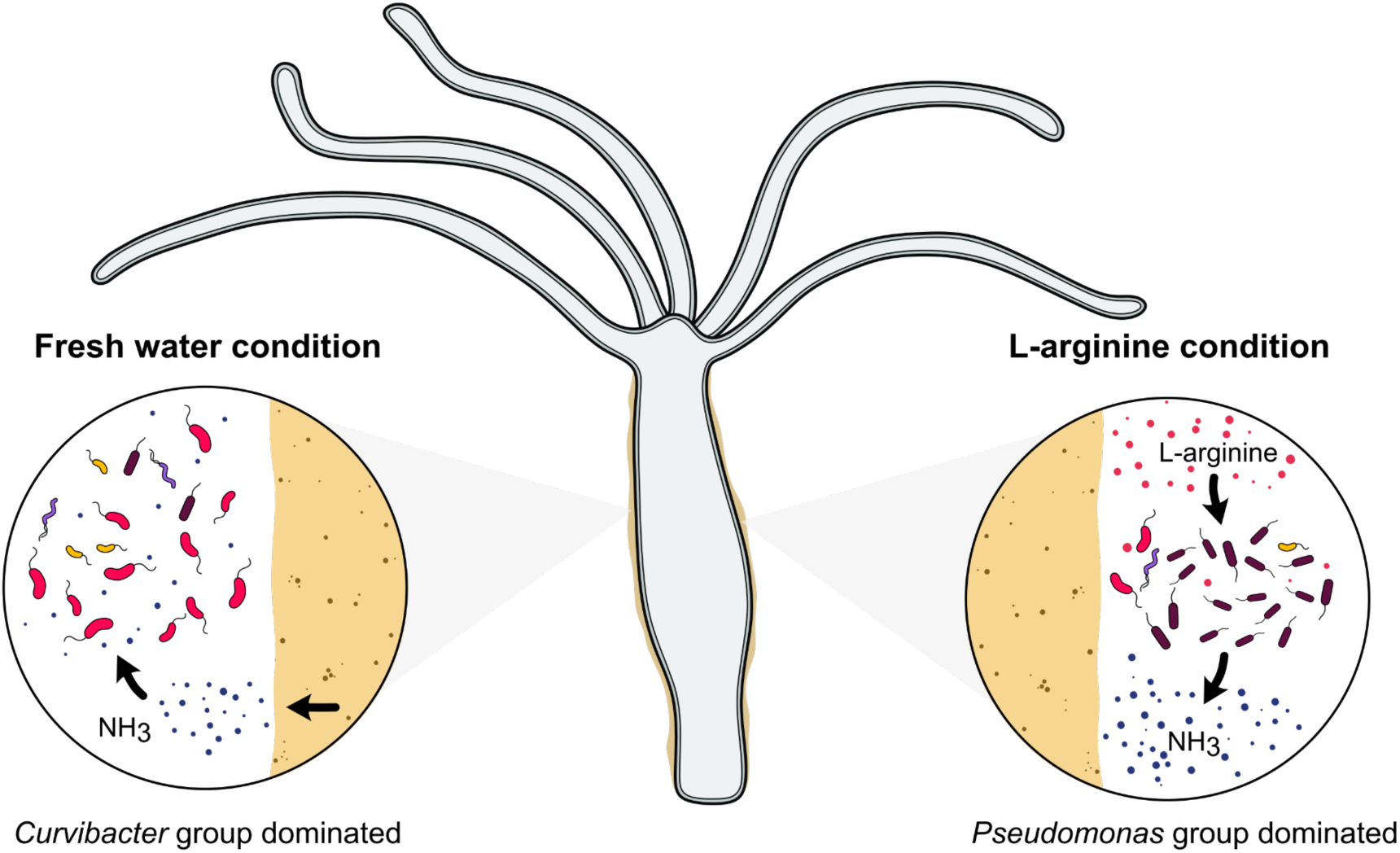
Summary of ammonia metabolism within the *Hydra*-microbe system under fresh water and L-arginine supplemented condition. Microbes in the *Curvibacter* group utilize ammonia as an electron donor, potentially cross-fed from the host in the fresh water condition, while under L-arginine supplement, *Pseudomonas* becomes dominant and shifts metabolism towards ammonia reduction.

Our study links amino acid availability, microbial metabolism, and host outcomes and thereby provides a simplified framework to interpret analogous processes in more complex systems such as the vertebrate gut. Controlled dietary intervention studies in humans showed rapid shifts in microbial gene abundance/expression and metabolic activity (21,71–73). Higher protein intake has been associated with enrichment of proteolytic taxa and increased microbial pathways involved in amino acid degradation and nitrogen metabolism (21,73,74). These functional changes lead to the increased production of ammonia and related metabolites, which have been linked to epithelial stress (75), increased renal toxicity (72), tumorigenesis (76,77) and worse outcome in dextran sodium sulfate (DSS) induced colitis (78). In contrast to the relatively buffered vertebrate gut microbiome, the *Hydra* system shows a tighter coupling between metabolite availability, community restructuring, and functional output. Nevertheless, the underlying ecological principle is conserved: shifts in resource availability can reprogram microbial metabolism and, consequently, host-microbe interactions.

With this study, we demonstrate a data-mining framework that extends 16S rDNA Biolog datasets to extract functional information. However, despite the robustness of our analysis, several limitations should be acknowledged. First, the identification of functional modules may include false positives, as microbes can carry certain functional genes without necessarily expressing or utilizing them under the tested conditions. Secondly, the 16S rDNA-genome linking step may assign multiple 16S sequences to the same genome due to the low specificity of 16S sequences and the limited sensitivity of genome assembly. Thirdly, our analysis is restricted to linkable 16S sequences, which, although they represent a large proportion of the community, do not cover it entirely. Finally, partial gene sets mapped to specific modules may lead to the apparent detection of functions not typically associated with microbes, such as plant- or animal-related modules. These observations reflect shared or analogous biochemical pathways rather than true cross-kingdom functions.

In summary, we systematically evaluated a wide range of single-metabolite perturbations to examine their effects on host-associated microbial communities. This work provides a comprehensive description of microbiome alterations and suggests potential mechanistic links between specific metabolites and community dynamics, helping to explain patterns of community assembly within the host-microbe context. Our findings highlight the critical roles of both microbe-microbe and host-microbe interactions in shaping overall community structure, underscoring the need to account for these factors in future studies. Further, we found that the *Hydra* microbial community composition is either unchanged or ends up in very similar compositions after perturbations (Supplementary Figure 13). This pattern is consistent with the idea that microbial communities assemble deterministically and occupy stable configurations, as proposed by previous theoretical and experimental studies (79–83). If the factors that triggered transitions between these states were known, microbial communities could potentially be steered through targeted perturbations. Our data indicate that in *Hydra*, such transitions are influenced by metabolic interactions among microbes, between microbes and the host, and with the environment. We therefore propose that tractable host-microbe model systems such as *Hydra* provide a valuable framework for identifying the drivers of state transitions and for developing predictive models of microbial community stability. Such insights will be essential for the rational control of microbiomes in medical and environmental applications.

## Materials and methods

### Taxonomic identification

To investigate microbial composition of the *Hydra vulgaris* AEP microbiome, 16S rDNA amplicons targeting the V1–V2 hypervariable regions were derived from microbial phenotypic microarray plates (BIOLOG) (12). These data provided taxonomic insights into the microbial community across ecologically diverse conditions in a total of 327 metabolite conditions (including control) across carbon (PM1 and PM2A), nitrogen (PM3B), and phosphorus and sulfur (PM4A) sources. Metabolite category information from Human Metabolome Database (HMDB) (84) was attached into each single-metabolite condition.

For the identification pipeline, amplicon sequencing data were initially assessed for quality using FastQC v0.12.1 (85) and summarized with MultiQC v1.17 (86). Taxonomic identification was performed using the DADA2 v1.26.0 (87). During the processing step, sequences were truncated to a length of 225 base pairs, and those with a phred quality score below 11 or more than 2 expected errors were removed. Subsequently, taxonomy was assigned using a Naive Bayes classifier trained on the SILVA nr99 database (v138.1_wSpecies) (88). To ensure a quality, microbial composition data underwent further filtering. Samples with a sequencing depth below 2,500 reads were excluded, based on rarefaction curves generated using the vegan v2.6.8 (89). Microbial taxa present at less than 1% abundance across all samples were classified as rare and excluded from downstream analyses. The quality check steps were conducted in Bash and the rest steps were done in R v4.4.1.

### DNA-extraction and metagenomic sequencing

For metagenomic analysis, 50 *Hydra vulgaris* polyps per sample were transferred into 1.5 ml microcentrifuge tubes. Residual *Hydra* medium was carefully removed and replaced with 300 µl phosphate-buffered saline (PBS). Samples were briefly vortexed, and 200 µl of the suspension was transferred to fresh tubes. Lysis was initiated by adding 22 µl 2M Tris-HCL (pH 8.5; 0.2M EDTA), 20 µl 10% (w/v) SDS, and 6 µl Proteinase K, followed by incubation at 37 °C for 20 min and 56 °C for 15 min.

Subsequently, 500 µl CTAB buffer (100 mM Tris-HCl, pH 8.0; 3 M NaCl; 20 mM EDTA; 3% [w/v] CTAB) was added, and samples were incubated at 65 °C for 15 min. An equal volume (500 µl) of chloroform:isoamyl alcohol (24:1) was added, and samples were centrifuged at 13,000 × g for 5 min at room temperature. The aqueous phase was transferred to a new tube and extracted with 700 µl chloroform:isoamyl alcohol, followed by centrifugation under the same conditions. This step was repeated until the aqueous phase was clear.

DNA was precipitated by adding 0.5 µl glycogen and 500 µl isopropanol, followed by overnight incubation at −20 °C. Samples were centrifuged at 13,000 × g for 20 min at 4 °C, and the supernatant was discarded. The resulting pellet was washed with 70% (w/v) ethanol, air-dried at room temperature, and resuspended in nuclease-free water.

Metagenomic libraries were prepared using the Nextera DNA Flex library preparation kit and sequenced on an Illumina NovaSeq 6000 platform (SP flow cell, 2 × 150 bp paired-end reads).

### Linking of 16s rDNA and microbial genomes

Since 16S rDNA sequences provide only taxonomic information, metagenome-assembled genomes (MAGs) reconstructed from shotgun metagenomic data, along with microbial genomes from the same host species, were associated with the 16S rDNA microbial taxa for functional information.

#### Reconstruction of metagenome assembled genome (MAG)

The MAGs of *Hydra vulgaris* AEP microbiome were reconstructed from shotgun metagenomic data. As the microbiome was from the same host species, it was assumed that microbial taxonomies were common and shared across two datasets.

The samples underwent a quality control and pre-processing pipeline to remove low-quality reads and technical contaminants. First, FastQC v0.12.1 (85) and MultiQC v1.17 (86) were used (86) initial quality assessments, and FastP v0.23.4 (90) was employed for trimming, with a minimum length cutoff of 30. Error base correction and low quality trimming was applied using default settings. Next, pre-processed reads were aligned to the *Hydra vulgaris* AEP genome (91) to exclude potential host genomic contaminants using BBmap v38.90 (92). Reads that did not align to the host genome were retained for subsequent MAG reconstruction. To construct MAGs, the filtered reads were processed through contig assembly, contig binning, and bin refinement steps using the metaWRAP v1.3.2 pipeline (93). Two assemblers, MEGAHIT v1.1.3 (94) and metaSPAdes v3.13.0 (95), were used in parallel to maximized assembled contigs possibly from diverse technical conditions. Assembled contigs were then binned using three separate binning tools, MetaBAT2 v2.12.1 (96), MaxBin2 v2.2.6 (97), and CONCOCT v1.0.0 (98), to capture a broader range of genome bins. Binned genomes were refined and consolidated using metaWRAP’s bin_refinement module, which merges bins from each binning tool and selects the best-quality bins. In this step, bins were retained if they met thresholds of ≥60% completeness and ≤40% contamination. Final bins from each sample, produced by both assemblers, were de-replicated using dRep v3.5.0 (99), with a threshold of ≥70% completeness and ≤20% contamination to generate a non-redundant, high-quality consensus set of MAGs. All steps were conducted in Bash and Snakemake v7.32.4 (100) was used to build analysis pipelines.

In addition to the reconstructed MAGs, microbial genomes associated with *Hydra vulgaris* AEP were included further for the next linking steps. These genomes included isolates from *Acidovorax*, *Duganella*, *Pseudomonas*, *Curvibacter*, *Pelomonas*, and *Undibacterium* (20), which were derived from monoculture experiments. For DNA isolation bacteria were grown in liquid R2A media (glucose 0.5 g/L, casamino acids 0.5 g/L, starch 0.5 g/L, proteose peptone 0.5 g/L, yeast extract 0.5 g/L, sodium pyruvate 0.3 g/L, K2HPO4 0.3 g/L, MgSO4 x 7 H2O 0.05 g/L) at 18°C to stationary phase and DNA was extracted using the DNAeasy Blood and Tissue Kit (Qiagen) following manufacturers instruction. Paired-end libraries were generated using the Illumina TruSeq LT kit (median fragment size: 402 bp), and mate-pair libraries were constructed with the Illumina Nextera Mate Pair Kit (insert size: 7.3 kb). Sequencing was performed on a MiSeq platform at the Biomolecular Resource Facility, The Australian National University, Canberra, Australia. Read quality filtering was done using Trimmomatic (leading 3, trailing 3, minlength 35nt, an overall crop of 250nt and a sliding window of 4:20) (101) and reads were assembled using SPAdes (102) (default settings). QUAST (103) was used for quality checks on the genomes.

#### Linking of 16s rDNA and microbial genomes

The linkage of 16S rDNA sequences to microbial genomes was conducted in two steps to maximize the coverage of 16S rDNA associations with the genome data. In the first step, we directly linked 16S rDNA to genomes using MarkerMAG v1.1.28 (104). MarkerMAG aligns shotgun metagenomic reads to both experimental 16s rDNA and microbial genomes, bridging them based on sequence identity and alignment depth. This approach effectively links 16S rDNA with genomes that are more complete and exhibit fewer mismatches. However, some genomes may lack complete 16S rDNA regions due to assembly gaps, preventing successful pairing between markers and genomes.

To increase linkage coverage, a second, indirect linkage step was implemented. In this step, we utilized three sets of predicted 16S rDNA sequences associated with genomes: (I) reconstructed 16S rDNA derived from shotgun metagenomic data using MATAM v1.5.3 (43), which were then linked to microbial genomes via MarkerMAG; (II) predicted 16S rDNA sequences from microbial genomes using barrnap (https://github.com/tseemann/barrnap); and (III) close reference 16S rDNA sequences of the microbial genomes from GTDBTk v2.4.0 (44,45). MATAM and barrnap functionalities embedded in MarkerMAG were utilized for this process. Next, all predicted 16S rDNA sequences, along with experimental 16S rDNA sequences, were compared through phylogenetic analysis. Initially multiple sequence alignment (MSA) was performed using MUSCLE v5.1 (105), and the most alignable regions in the MSA were selected using GBLOCKS v0.91b (106). A phylogenetic tree was then reconstructed with FastTree (107), and phylogenetic distances were computed from the resulting tree using ape v5.8 (108) within R v4.4.1. Pairs between predicted and experimental 16S rDNA sequences with a phylogenetic distance of ≤5% were retained for further analysis, while those exceeding this threshold were excluded from further analysis. Apart from phylogenetic distance calculation, the other steps were conducted in Bash.

Since limitation in taxonomic identification of the 16s rDNA and sensitivity of genome assembly from pooled reads, the detailed information of microbial sequences might not be able to explain species-or strain-level variability. Thus, a microbial genome was allowed to link to one or more amplicon sequences.

### Functional annotation and quantitation

The genomes linked to 16S rDNA sequences were further annotated to determine their functional potential. Metabolic capacity was quantified for each genome to assess its potential ecological role, and this information was subsequently used to estimate the overall metabolic capacity of each microbial community by incorporating microbial abundance.

First, functional annotation of the microbial genomes was performed using eggNOG-mapper v2.1.18 (109) which utilizes Diamond v2.1.9 (110) for sequence alignment and Prodigal v2.6.3 (111) for gene prediction. From this annotation, KEGG Orthology (KO) and KEGG Module (KM) terms were derived from the KEGG database to enable functional quantification (112). Next, to quantify functional potential for each genome, KEGG module coverage 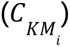 and KEGG module abundance 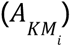 were calculated according to Equations 1 and 2.

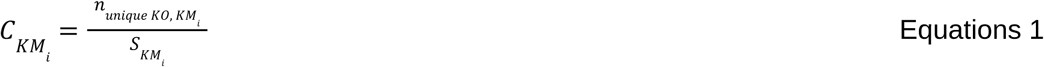

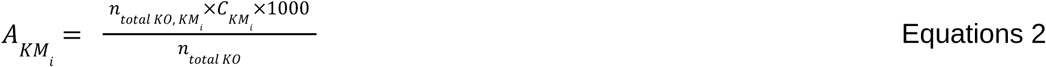

where:

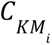: Coverage of KEGG Module for each genome.

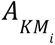: Abundance of KEGG Module for each genome.

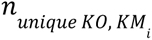: Nonredundant (unique) count of KEGG Orthologies in KEGG Module.

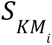: Total number of KEGG orthologs in KEGG Module.

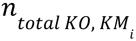: Number of KEGG orthologs in KEGG Module found in each genome.

*n*_*total* *KO*_: Total count of KEGG Orthologies across all KEGG Modules.

Following genome-level quantification, these metrics were integrated with microbial abundance data to calculate the community level module abundance (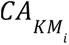; Equation 3).

This approach allowed us to link functional annotations to community-level metabolic potential. The functional annotation steps were conducted in Bash and the calculations were done in R v4.4.1.

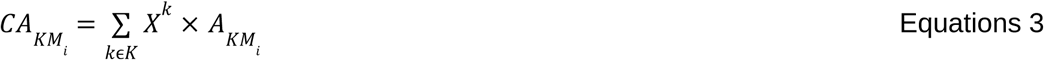

where:

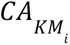: Abundance of KEGG Module for each microbial community.

*X^k^*: Relative abundance of microbial k in a microbial community.

### Analysis of microbial taxonomy and function

All analysis steps below were conducted in R v4.4.1.

#### α- and β-diversity analysis

α-diversity was assessed using the richness, evenness, and Shannon index, calculated using vegan v2.6.8 (89). To evaluate the impact of metabolite supplementation at both the subclass and individual metabolite levels on α-diversity, linear mixed models were fitted using the lmer function from the lme4 v1.1.35.5 (113). In these models, the Biolog plate was included as a random effect to account for potential variability in initial microbial composition across plates, assuming that metabolite effects on α-diversity are consistent despite plate-to-plate variation. Next, for pairwise comparisons, the same linear mixed model approach was used when the metabolite condition appeared in more than one Biolog plate. For metabolite conditions unique to a single plate, linear models were fitted using the lm function from stats v4.4.1. β-diversity was calculated using Bray–Curtis dissimilarity index, betadisper function was used to assess group homogeneity. Pairwise comparisons between control and non-control conditions were performed using PERMANOVA with adonis2 function. All steps were done using the vegan package. The significant metabolite conditions either from the α or β-diversity analysis further underwent metabolite category enrichment analysis using an enricher command from clusterProfiler v4.14.4 (114). Multiple testing correction was performed using the method by Benjamini–Hochberg for all analyses.

#### Genome-level functional coverage analysis

Functional coverage profiles were analyzed by clustering similar functional profiles and filtering the modules which show high variation across groups. First, a similarity matrix was generated using the Jaccard index, calculated with vegan v2.6.8 (89). Hierarchical clustering was then applied to group genomes based on their functional profiles. The optimal number of clusters was determined using the within-cluster sum of squares approach implemented in the factoextra v1.0.7. Ordination of Jaccard similarity on the functional profiles was conducted using Non-metric Multidimensional Scaling (NMDS) by metaMDS function from vegan. To identify functionally relevant KEGG modules, the Kruskal–Wallis test was applied to each module to assess differences in functional coverage across clusters. Modules with statistically significant differences (corrected p < 0.05, Benjamini–Hochberg correction) were further filtered by Gini index using DescTools v0.99.58 (115). Only modules with a Gini index larger than 0.5 and a standard deviation greater than 0.1 were retained for visualization.

#### Community-level functional abundance analysis

We performed microbial community differential functional abundance analysis by comparing community module abundance of control conditions to the single metabolite supplemented condition. The linear mixed model was used when the metabolite condition appeared in more than one Biolog plate while (plate as random factor), for metabolite conditions unique to a single plate, linear models were fitted using the lm function from stats v4.4.1. P values were corrected using the Benjamini–Hochberg, applied within each plate based on the number of metabolite conditions tested.

#### Taxonomy-group differential abundance analysis

We performed differential abundance analysis based on pooled 16S sequence counts according to the functional clusters. For each sample, the 16S sequence counts of all taxa in the same functional cluster were added together. The group level count was then used as the input for ANCOMBC v2.8.0 (116) for comparison against the control condition. Biolog plates were used as a random effect in the linear model to correct for the bath effect. P values were corrected using the Benjamini–Hochberg.

### Experimental validation

#### In vitro assays

We first assessed the ability of *Pseudomonas* to generate ammonia from L-arginine, and the capacity of representatives of the *Curvibacter* group (*Curvibacter* and *Limnobacter*) to reduce environmental ammonia concentrations.

Bacterial strains (*Pseudomonas*, *Curvibacter*, and *Limnobacter*) were cultured in R2A medium (ROTH®) at 18 °C under shaking conditions. Exponentially growing overnight cultures were centrifuged at 6,000 × g for 5 min, and supernatants were removed. Cell pellets were resuspended in sterile nutrient-free mineral base medium (S-medium) and centrifuged again under the same conditions. Following removal of the supernatant, pellets were resuspended in fresh sterile S-medium. Thus, a total of two consecutive washing steps were performed to remove residual culture medium. Final cell densities were adjusted to OD_600_ = 0.2.

To mimic metabolite perturbation conditions identified in the computational analysis, bacteria were tested either in monoculture or in co-culture (*Pseudomonas* + *Curvibacter*; *Pseudomonas* + *Limnobacter*). For each treatment, 50 µl of bacterial suspension was added to 3 ml of one of the following conditions: (i) L-arginine (0.3 mg/ml), (ii) ammonia (5.86 µg/ml), or (iii) S-Medium (control). In co-culture treatments, 50 µl of each strain was added. Monocultures were volume-adjusted with 50 µl sterile S-medium.

All treatments were performed with five biological replicates (n = 5). Following 24 h incubation at 18°C, ammonia concentrations were quantified using the Ammonia Assay Kit (Sigma-Aldrich), according to the manufacturer’s instructions.

#### In vivo assays in *Hydra*

To assess whether the observed metabolic interactions translate to the host context, we performed recolonization experiments using germ-free *Hydra vulgaris* AEP polyps.

Germ-free polyps were generated following established protocols (10). Briefly, animals were washed 12 h after feeding and incubated in antibiotic solution (50 mg/ml each of ampicillin, rifampicin, spectinomycin, streptomycin, and neomycin in S-medium) at 18 °C in the dark. The solution was replaced every third day. After one week, polyps were transferred to sterile S-medium for 2 days to remove residual antibiotics.

For recolonization, germ-free polyps were washed and transferred to sterile S-medium (15 polyps per 50 ml). Bacterial suspensions (OD_600_ = 0.2) were added at 50 µl per strain. Co-colonization treatments received 50 µl of each strain, while monocultures were volume-adjusted with sterile S-medium.

After 2 days of recolonization, polyps were washed and transferred to 96-well plates. To replicate metabolite perturbation conditions, polyps were exposed to (i) L-arginine (0.3 mg/ml), (ii) ammonia (5.86 µg/ml), or (iii) S-medium (control), in a final volume of 200 µl per well. All treatments were performed with five biological replicates (n = 5).

After 24 h, ammonia concentrations were measured using the Ammonia Assay Kit (Sigma-Aldrich).

For statistical analysis, we conducted pairwise comparison between microbial treatments for each host, and environmental perturbing conditions using t-test and adjusted p-values with the Benjamini-Hochberg correction.

## Data Availability

All scripts for data analysis are available at github.com/nsrinak/MetPerturb-HydraMicrobe. The shotgun metagenomic data generated in this study are available in the NCBI BioProject database under accession number PRJNA1463651. All bacterial genomes reconstructed in this study are available at zenodo (https://doi.org/10.5281/zenodo.20084193).

## Author Contributions

NS - Methodology, software, formal analysis, investigation, data curation, writing original draft, review and editing, visualization

TL - Conceptualization, Methodology, validation, formal analysis, investigation, data curation, review and editing, project administration

LU - Investigation, data curation, review and editing

SF - Resources, review and editing, project administration, funding acquisition

CK - Review and editing, project administration, funding acquisition

JT - Conceptualization, investigation, resources, writing original draft, review and editing, supervision, project administration, funding acquisition

## Acknowledgments

We thank Tal Dagan, Robin Koch and Carina Knappe for help with the assembly of genomes from bacterial isolates. We are thankful to Claudia Taubenheim for critically reading and discussing the manuscript with us. We acknowledge support by the German Research Foundation within the framework of the Excellence Cluster “Precision Medicine in Chronic Inflammation” (project code EXC2167), the project ExoMod (project code KA 3541/20), and the research group miTarget (project code FOR5042) to C.K. and individual funding (TA1699.2, project code 525705073) to J.T.. Moreover, we acknowledge support by the German Ministry for Education and Research within the scope of e:Med iTREAT (project code 01ZX1902A) to C.K.. T.L., S.F. and C.K. acknowledge funding from the German Research Foundation within the framework of Collaborative Research Center 1182 of DFG Project-ID 261376515–SFB 1182, “Origin and Function of Metaorganisms”. S.F was supported by the German Research Foundation via project FR 3041/3-1. In addition, we acknowledge the usage of LLMs in the writing process of the manuscript. The LLMs were used to rephrase, shorten and increase readability of the original text. All text generated with these models was thoroughly checked for correctness and edited where needed to express our original ideas.

## Supplementary data

Supplementary File 1: Heatmap of community functional abundance changes

## Supplementary figure

**S.Fig.1.**
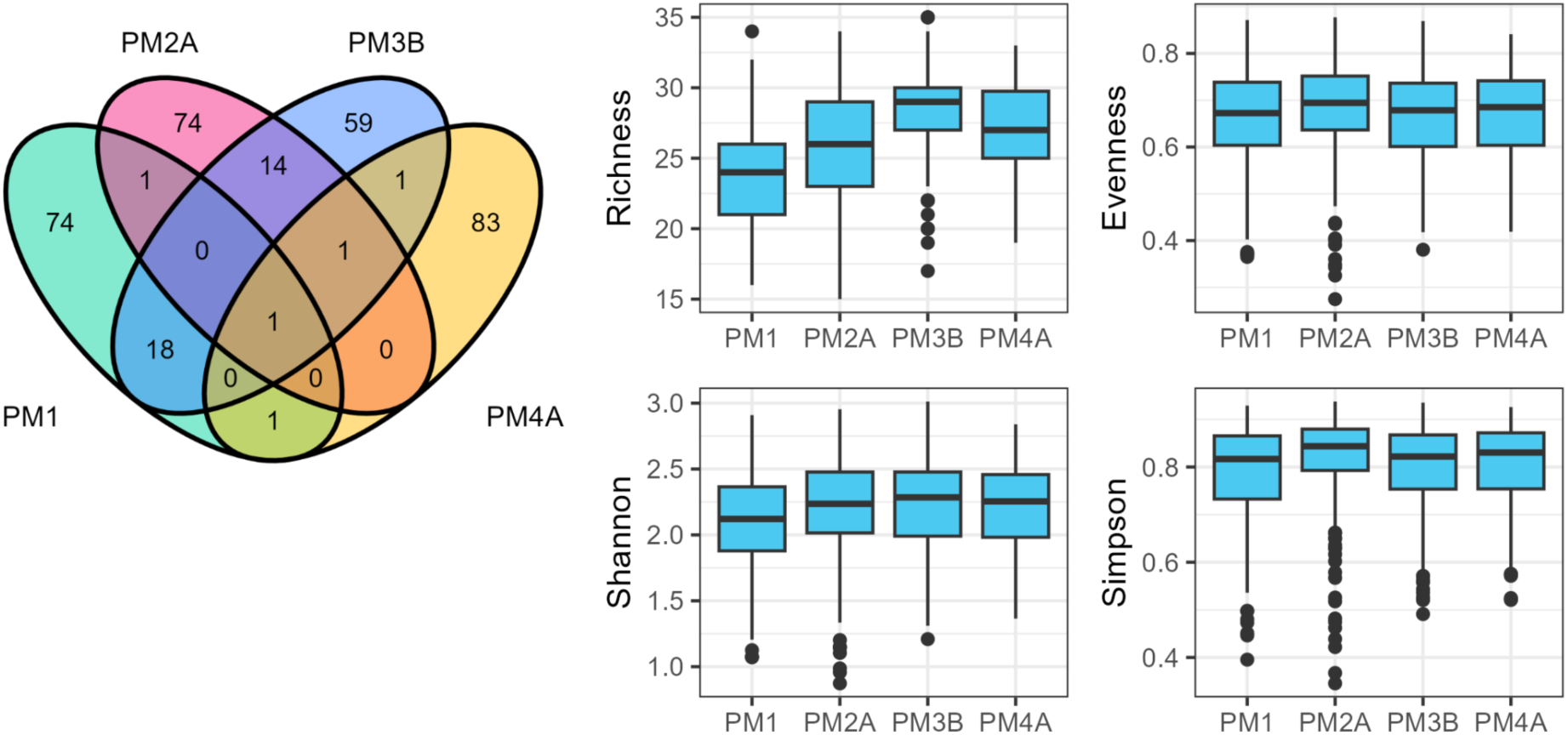
Venn diagram of metabolite conditions in different Biolog plates, PM1, PM2A, PM3B, and PM4A. Boxplots indicate microbial α-diversity in different Biolog plates.

**S.Fig.2.**
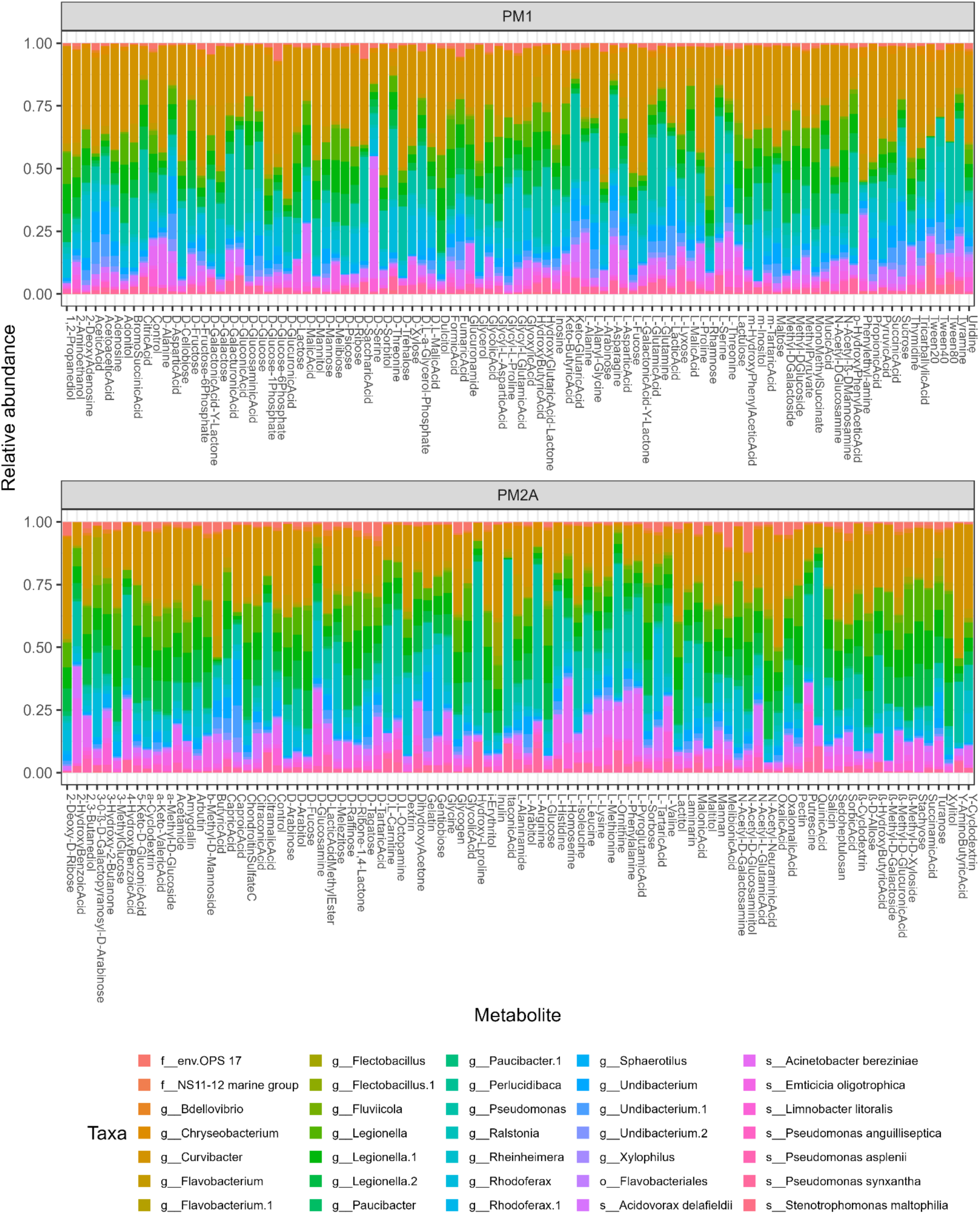
Relative abundance of microbial communities after metabolite perturbation in PM1 and PM2A Biolog plates.

**S.Fig.3.**
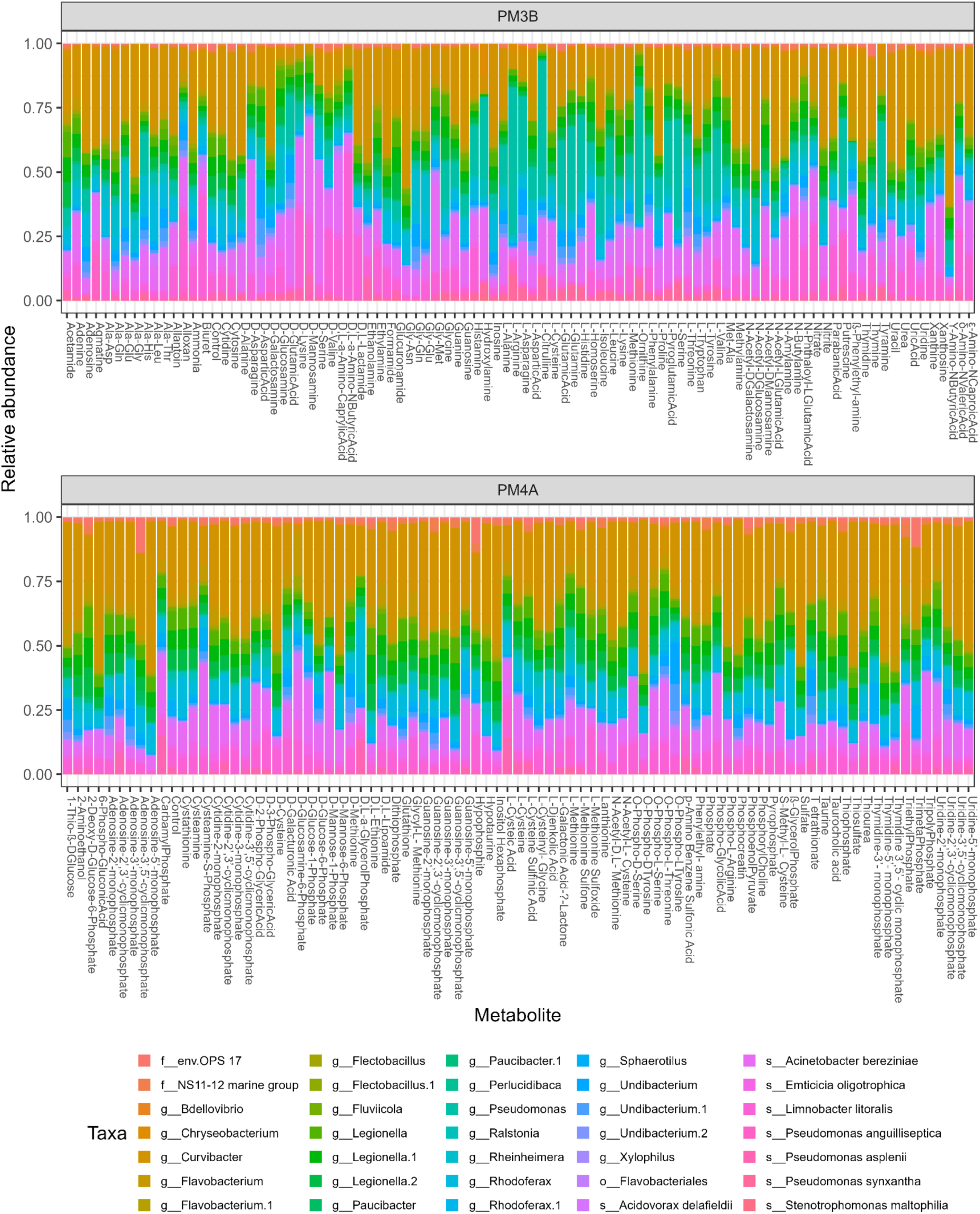
Relative abundance of microbial communities after metabolite perturbation in PM3B and PM4A Biolog plates.

**S.Fig.4.**
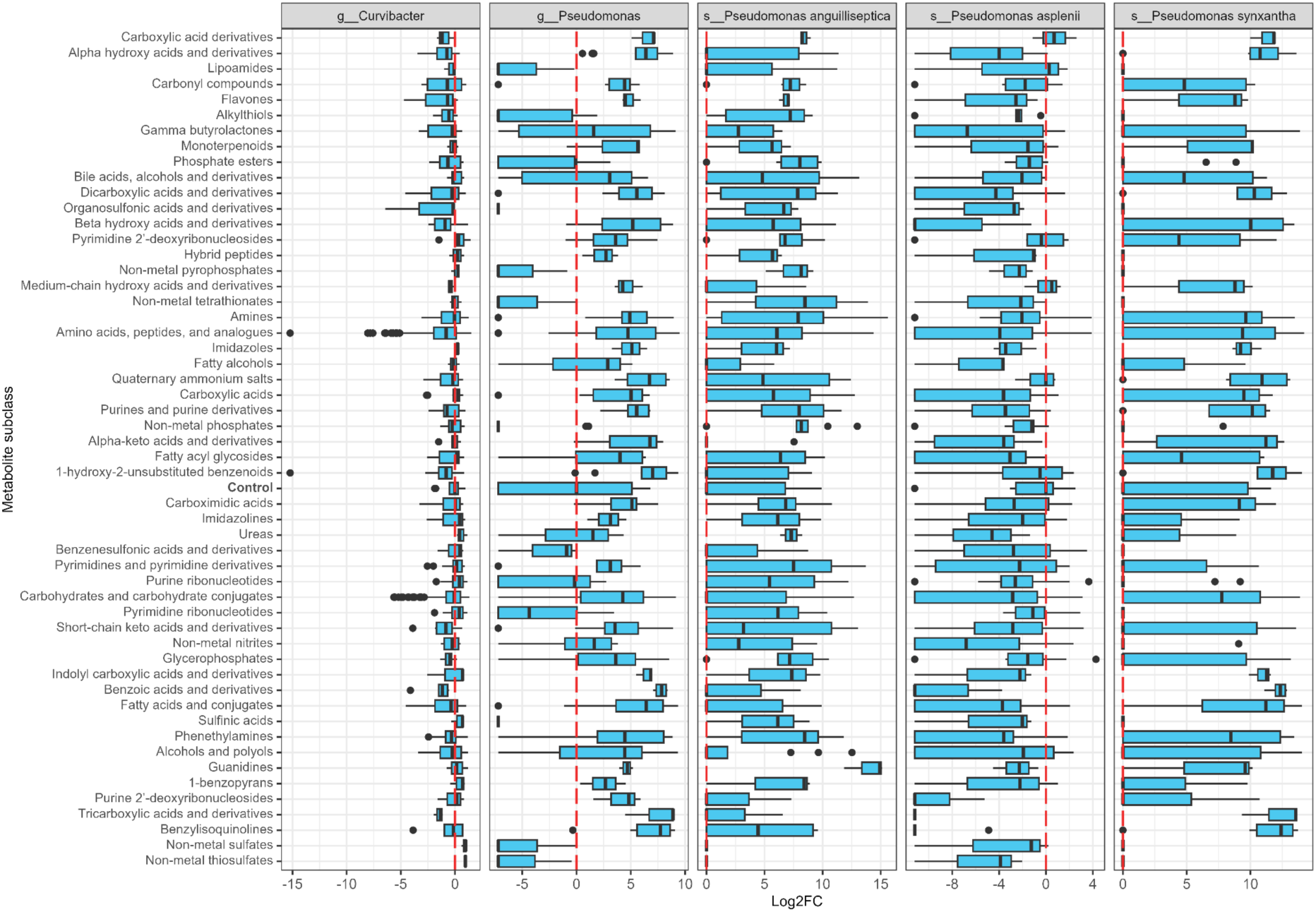
Log 2-fold change of relative abundance of *Curvibacter* and *Pseudomonas* species between testing and control conditions at metabolite subclass level.

**S.Fig.5.**
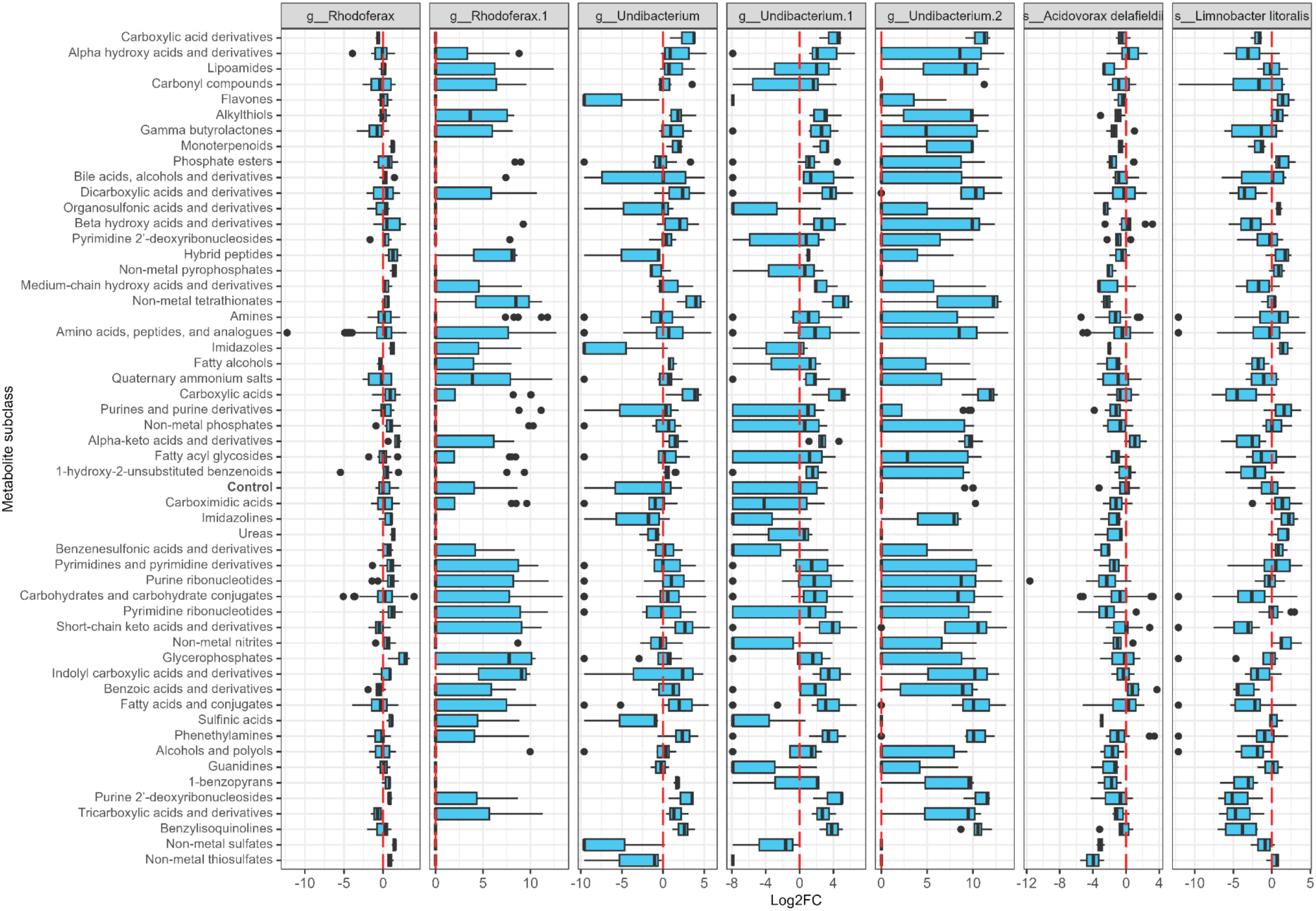
Log 2-fold change of relative abundance of high prevalent microbes between testing and control conditions at metabolite subclass level.

**S.Fig.6.**
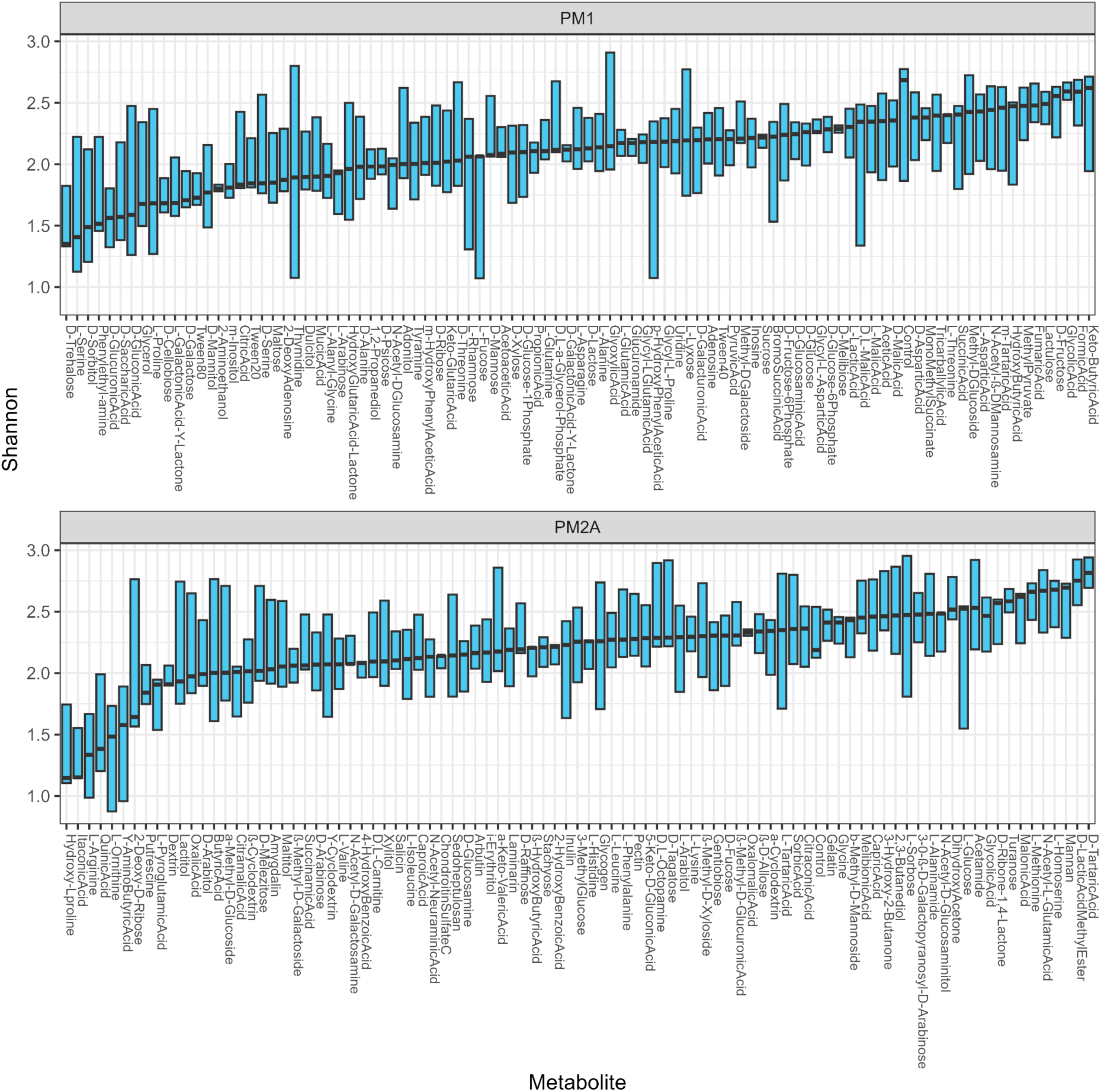
Shannon α-diversity of microbial communities after metabolite perturbation in PM1 and PM2A Biolog plates.

**S.Fig.7.**
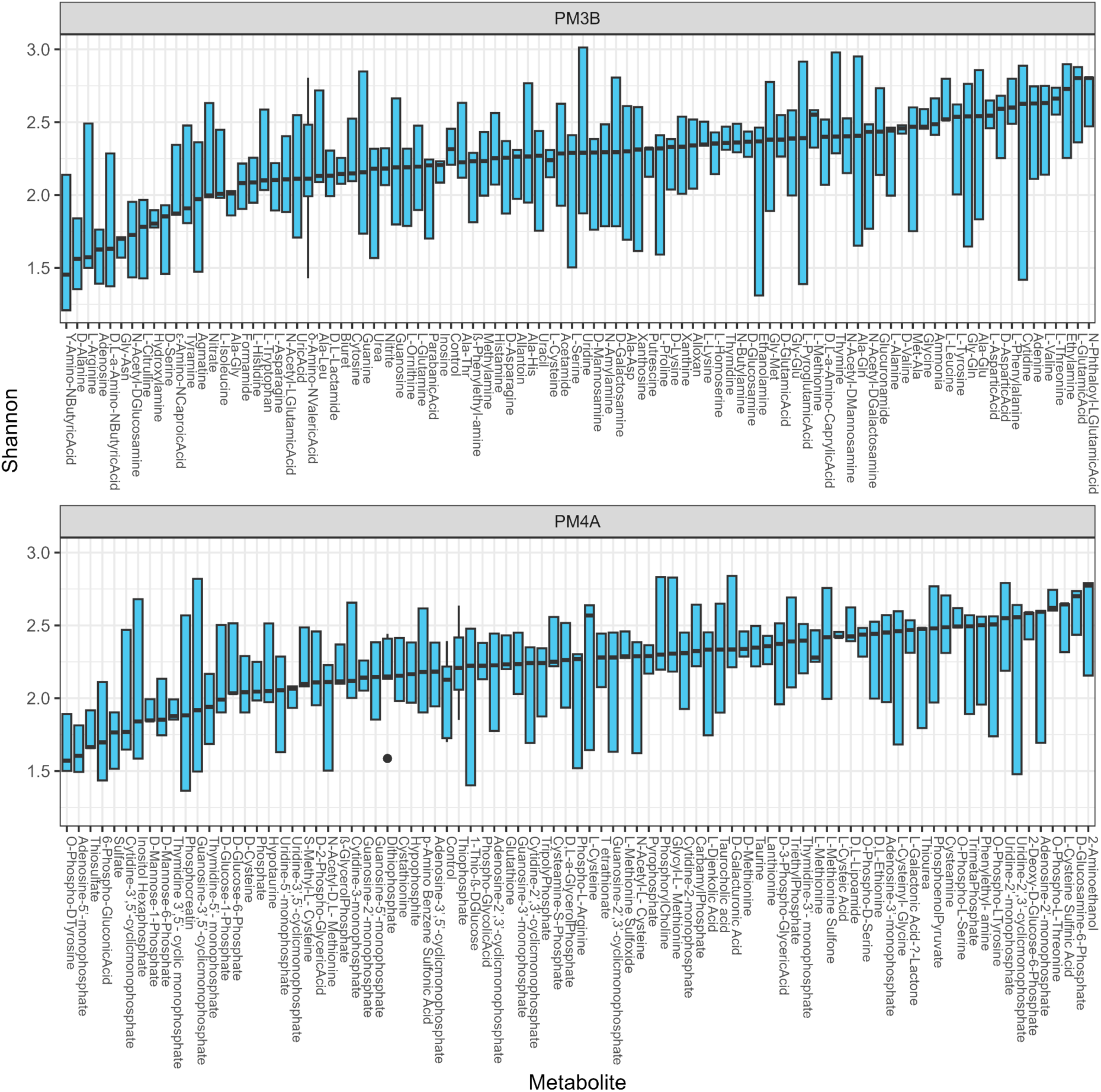
Shannon α-diversity of microbial communities after metabolite perturbation in PM3B and PM4A Biolog plates.

**S.Fig.8.**
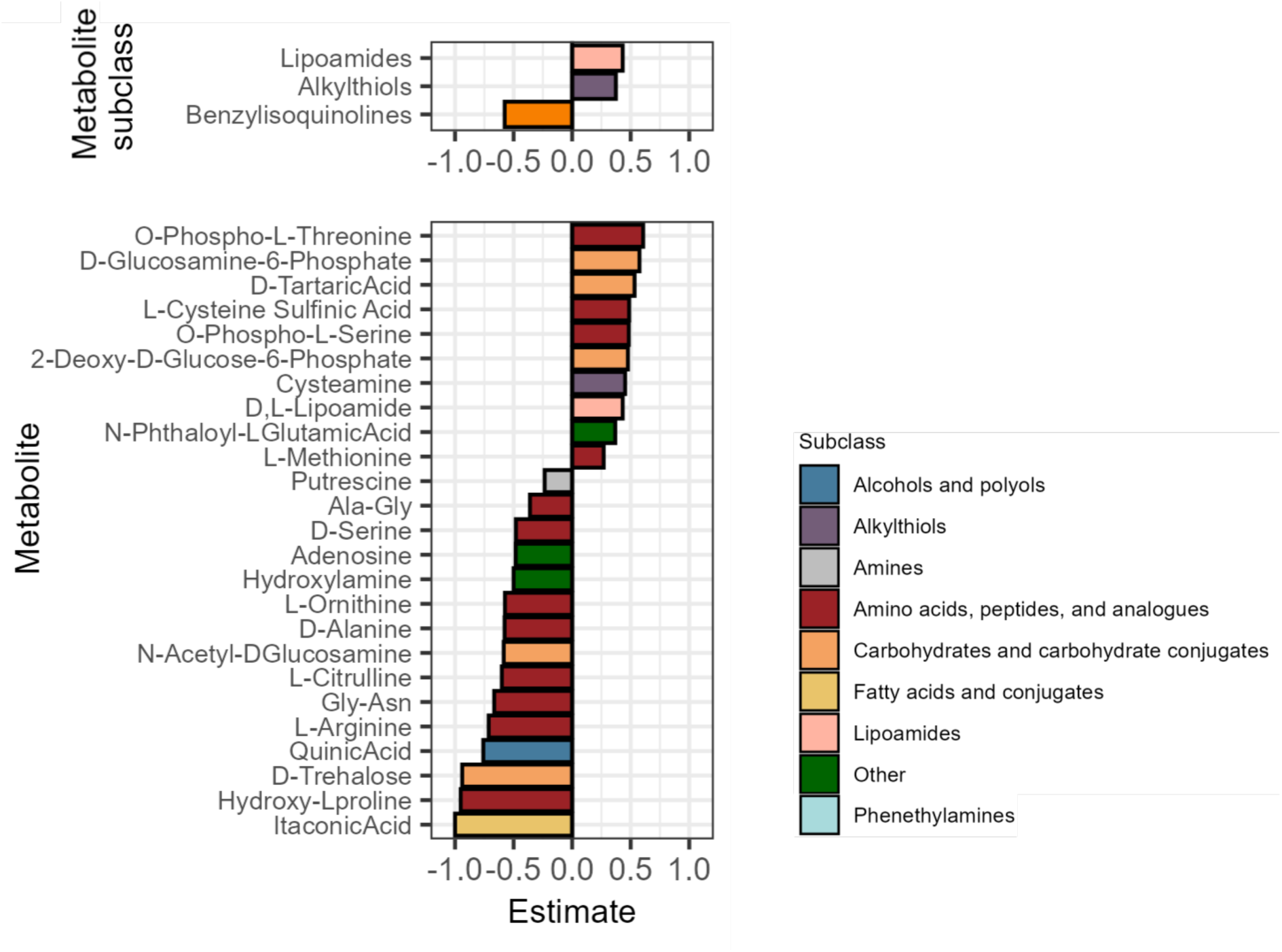
α-diversity changes in different metabolite-perturbed conditions. Estimated coefficients from linear mixed models show differences in Shannon index between each metabolite subclass (upper) or individual metabolite (lower) and the control condition (p < 0.05).

**S.Fig.9.**
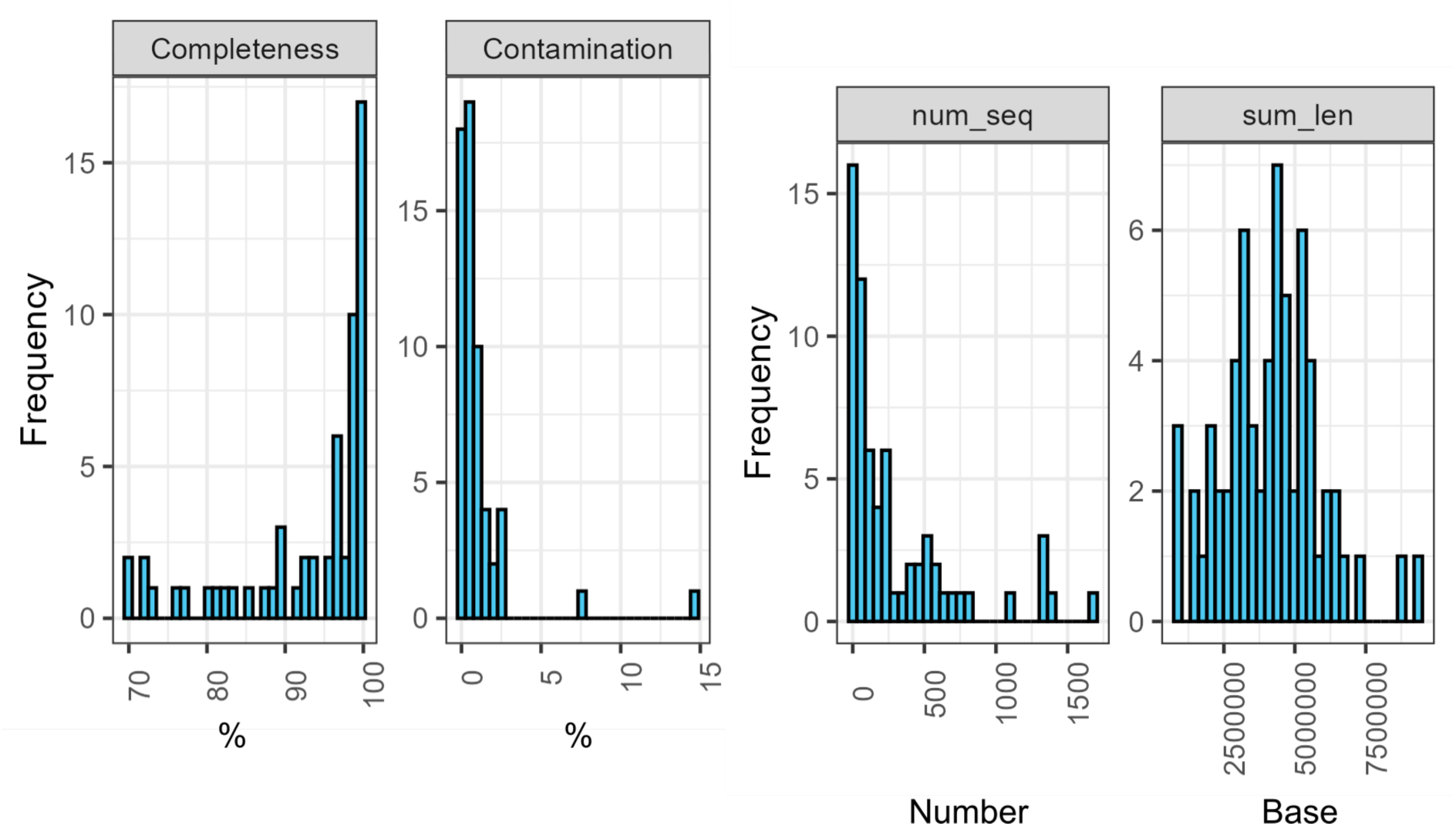
Completeness, contamination, number of sequences (num_seq), and total length (sum_len) of derived metagenome-assembled genomes (MAGs).

**S.Fig.10.**
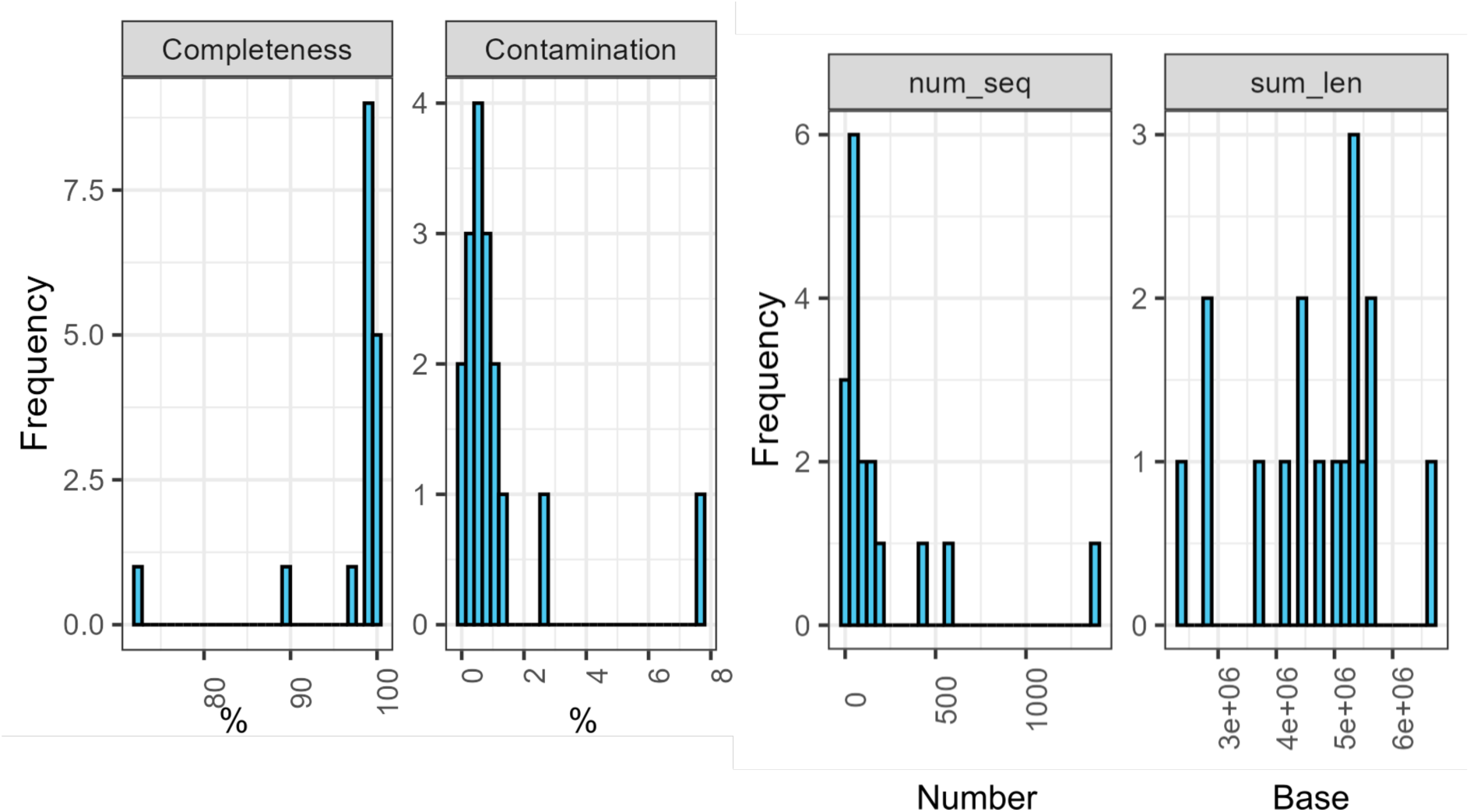
Completeness, contamination, number of sequences (num_seq), and total length (sum_len) of selected microbial genomes.

**S.Fig.11.**
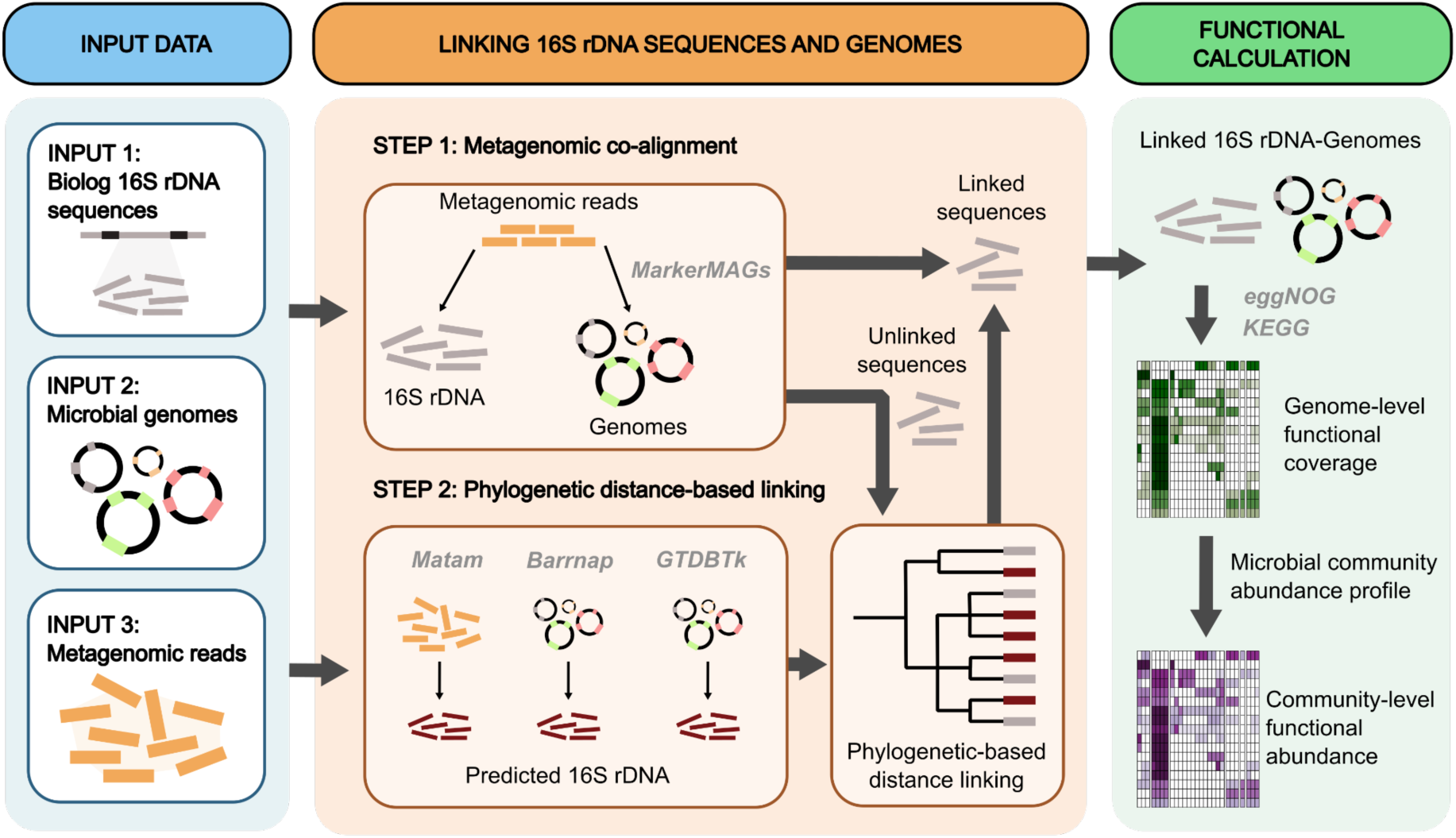
Workflow of 16s rDNA and microbial genomes linking approach.

**S.Fig.12.**
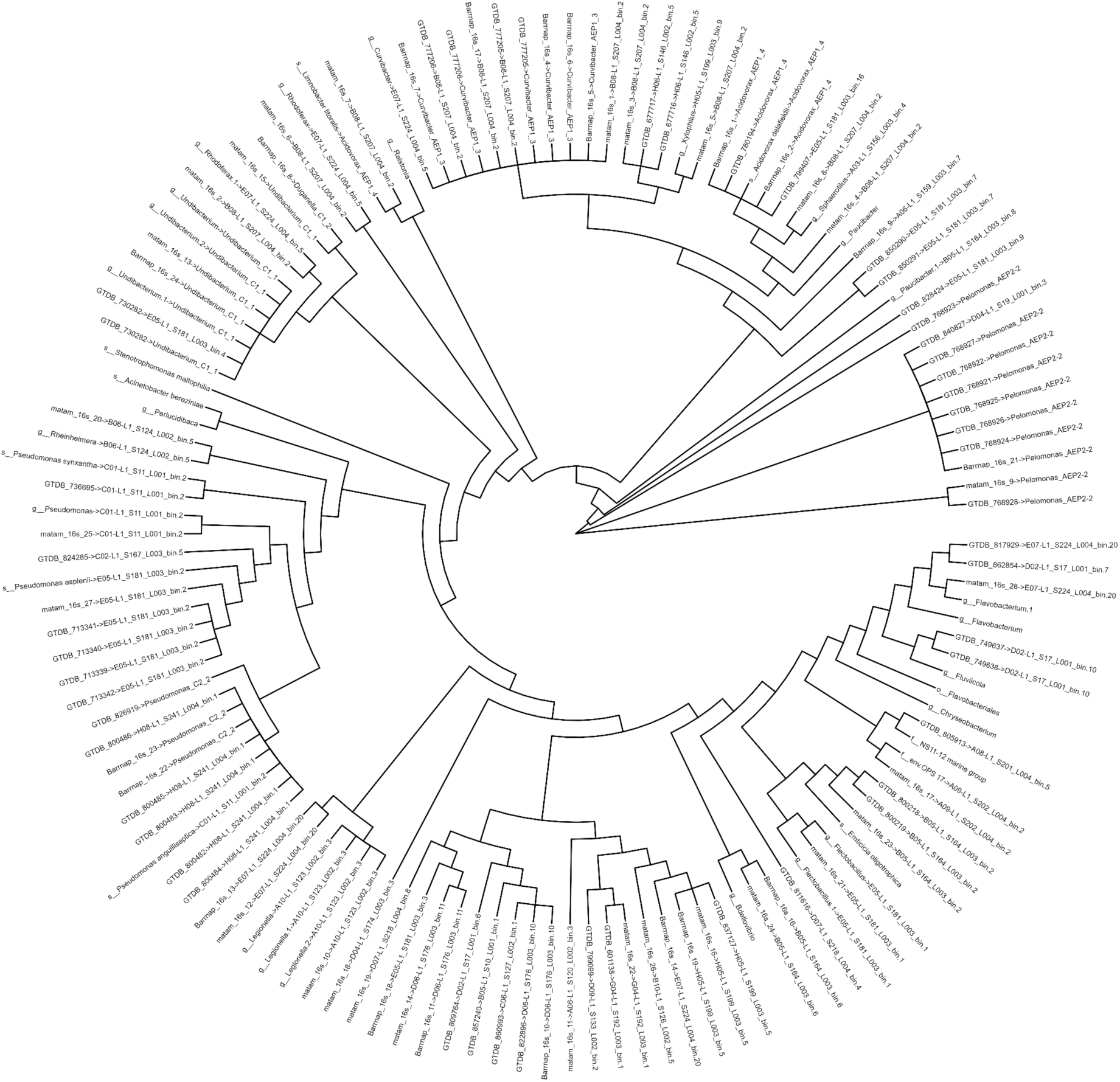
Phylogenetic tree of Biolog 16S rDNA and 16S rDNA from predictions: MATAM, barrnap, and GTDBTk. In tip labels, the arrows point from 16s or predicted 16s rDNA sequences to their linked genomes.

**S.Fig.13.**
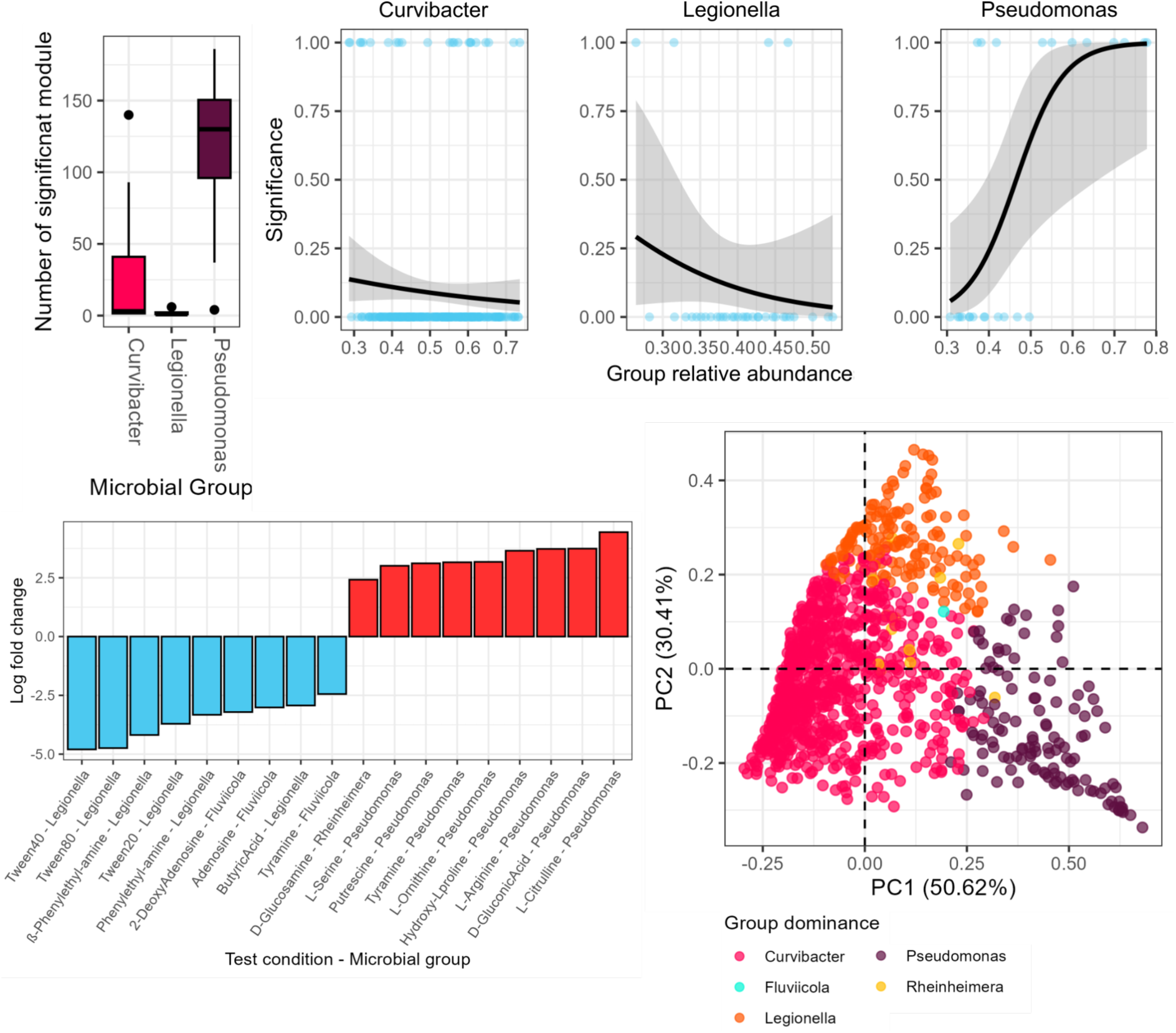
Microbial community dominance and functional change. The boxplot shows the number of significantly changed modules on each microbial group dominance. The dot plots show logistic associations between the relative abundances of microbial groups, i.e., *Curvibacter*, *Legionella*, and *Pseudomonas*, and significantly different functions. Binary values in dot plots, 0 and 1 denote the absence of significant functions or the presence of at least one significant function in a sample, respectively. The box plot shows log 2-fold change of microbial group abundance from ANCOM-BC between testing and control conditions at metabolite level (FDR < 0.05). Principal coordination analysis (PCoA) of Bray Curtis dissimilarity on microbial group relative abundance. The data points were labeled by the microbial group dominance in each sample.

**S.Fig.14.**
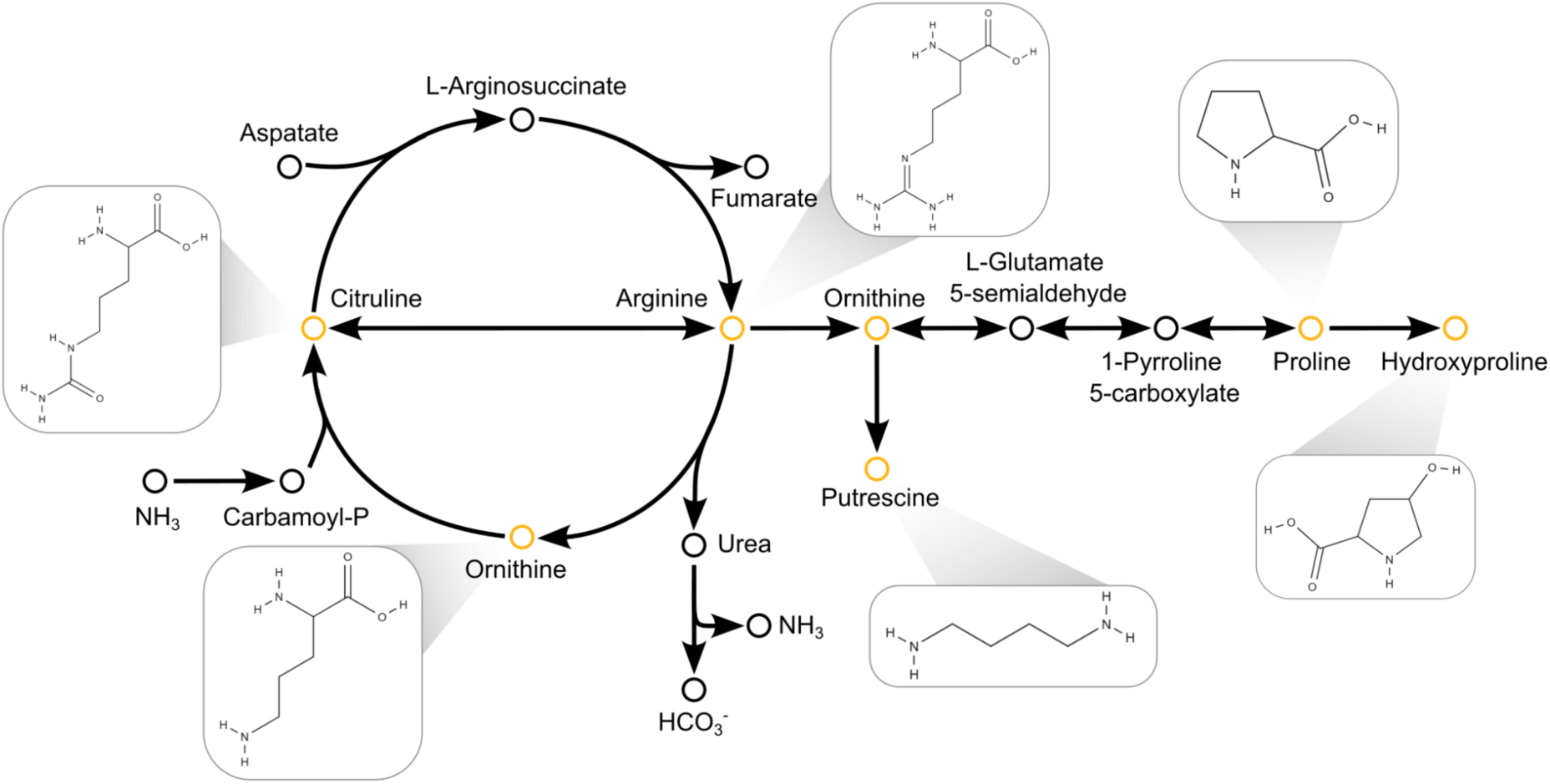
Metabolic reactions of L-arginine-related compounds. Orange nodes indicate the significant L-arginine-related compounds in this study. Chemical structures were generated using MolView (https://molview.org).

**S.Fig.15.**
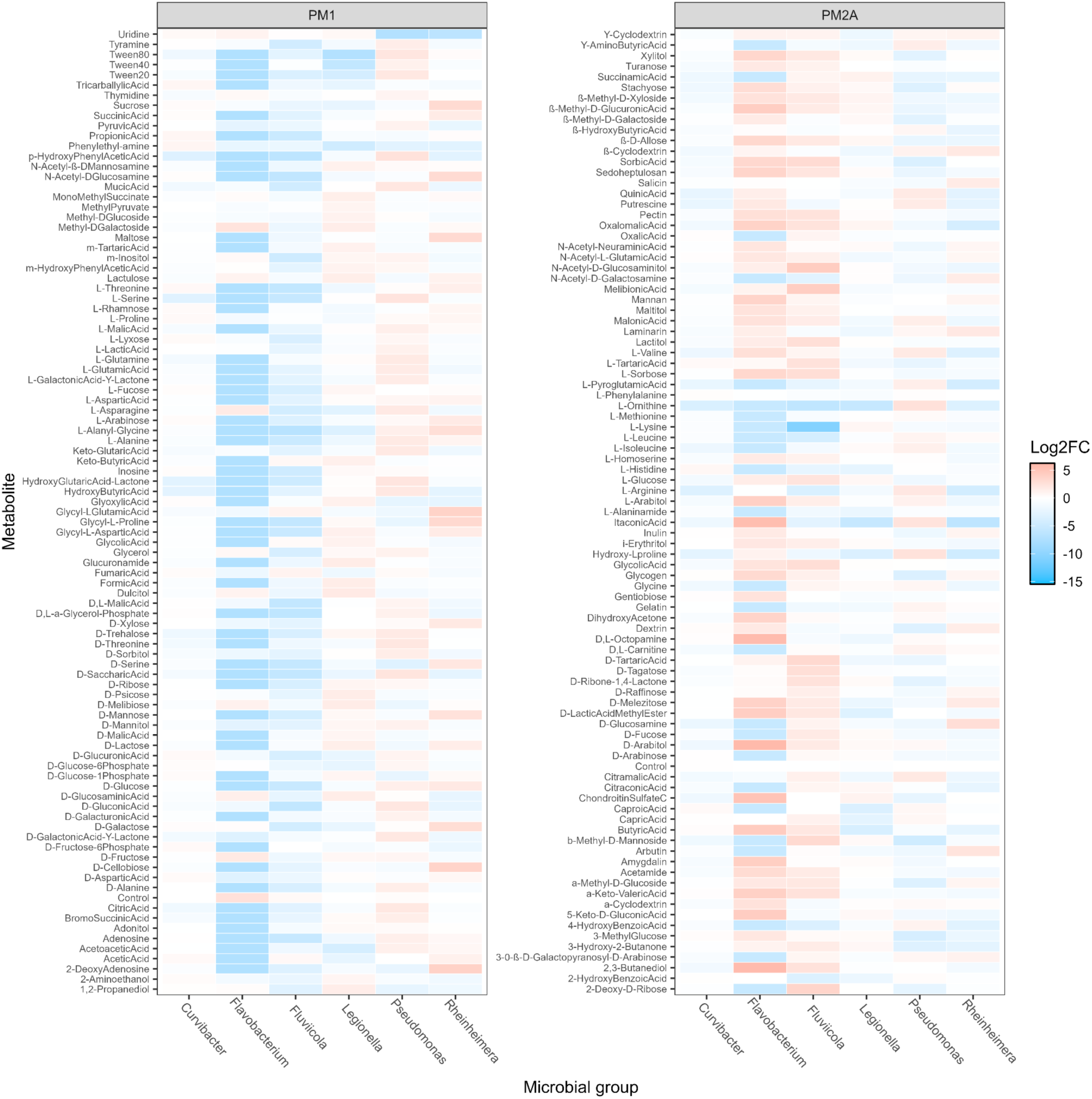
Log 2-fold change of relative abundance of microbial group between testing and control conditions at metabolite level in PM1 and PM2A Biolog plates.

**S.Fig.16.**
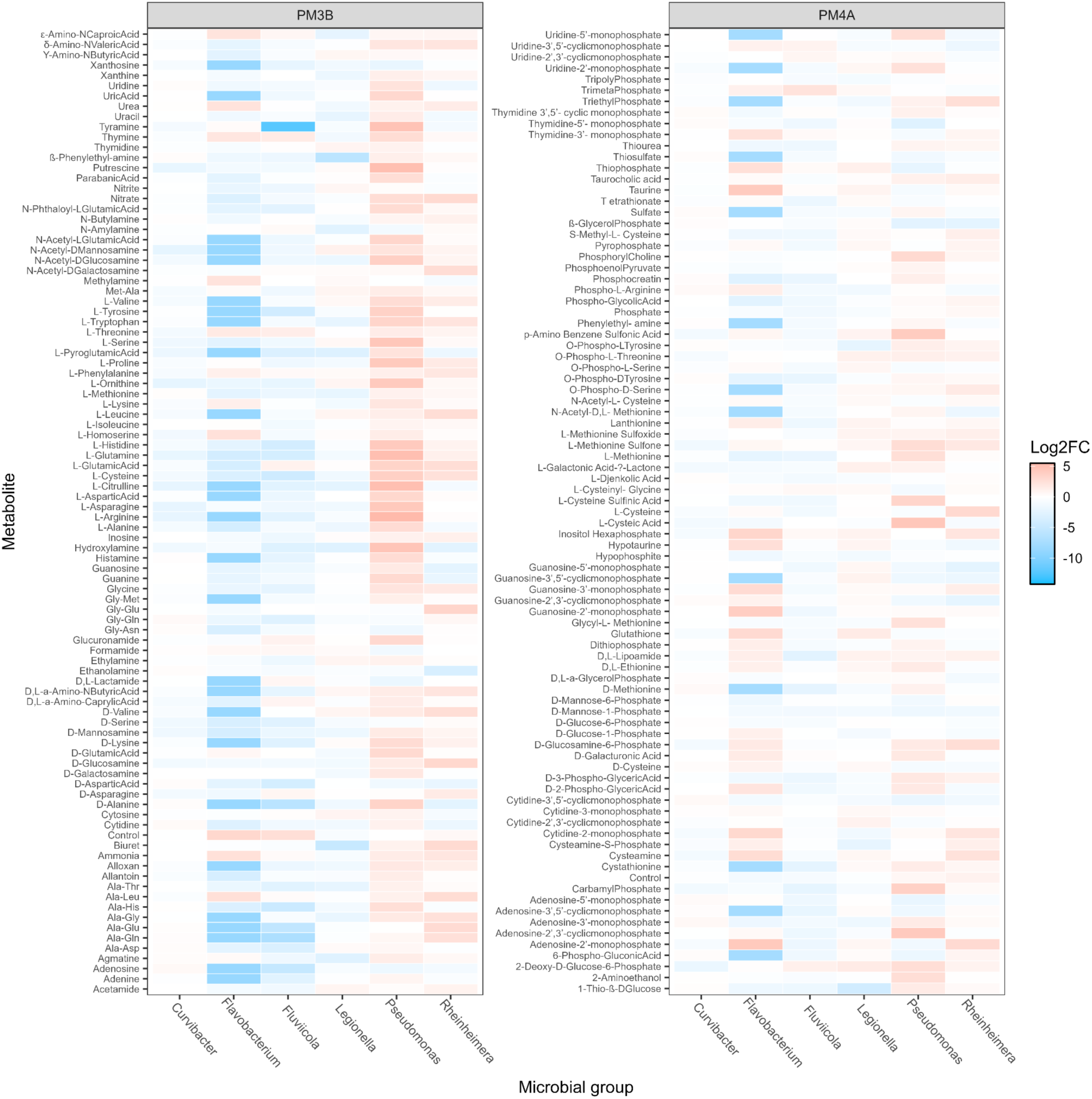
Log 2-fold change of relative abundance of microbial group between testing and control conditions at metabolite level in PM3B and PM4A Biolog plates.

**S.Fig.17.**
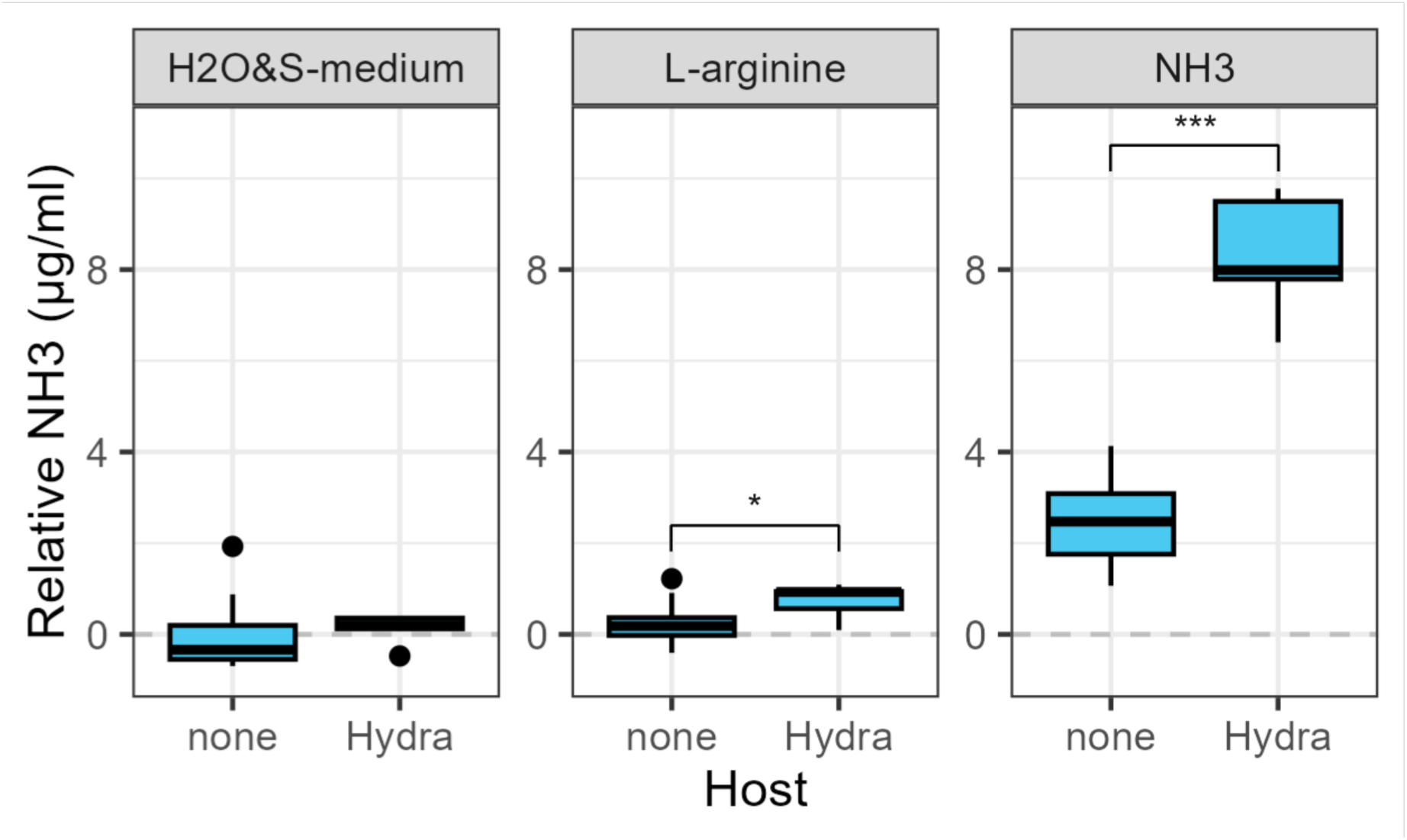
Ammonia concentration comparison in conditions with germfree *Hydra* and without *Hydra*. The comparisons of ammonia concentration were conducted in a normal condition (H_2_O in none host and S-medium in GF host), L-arginine, and NH_3_ supplemented conditions using t-test. Point and asterisk denote significance levels: P value < 0.10 (.), < 0.05 (*), < 0.001 (**), and < 0.0001 (***).

**S.Fig.18.**
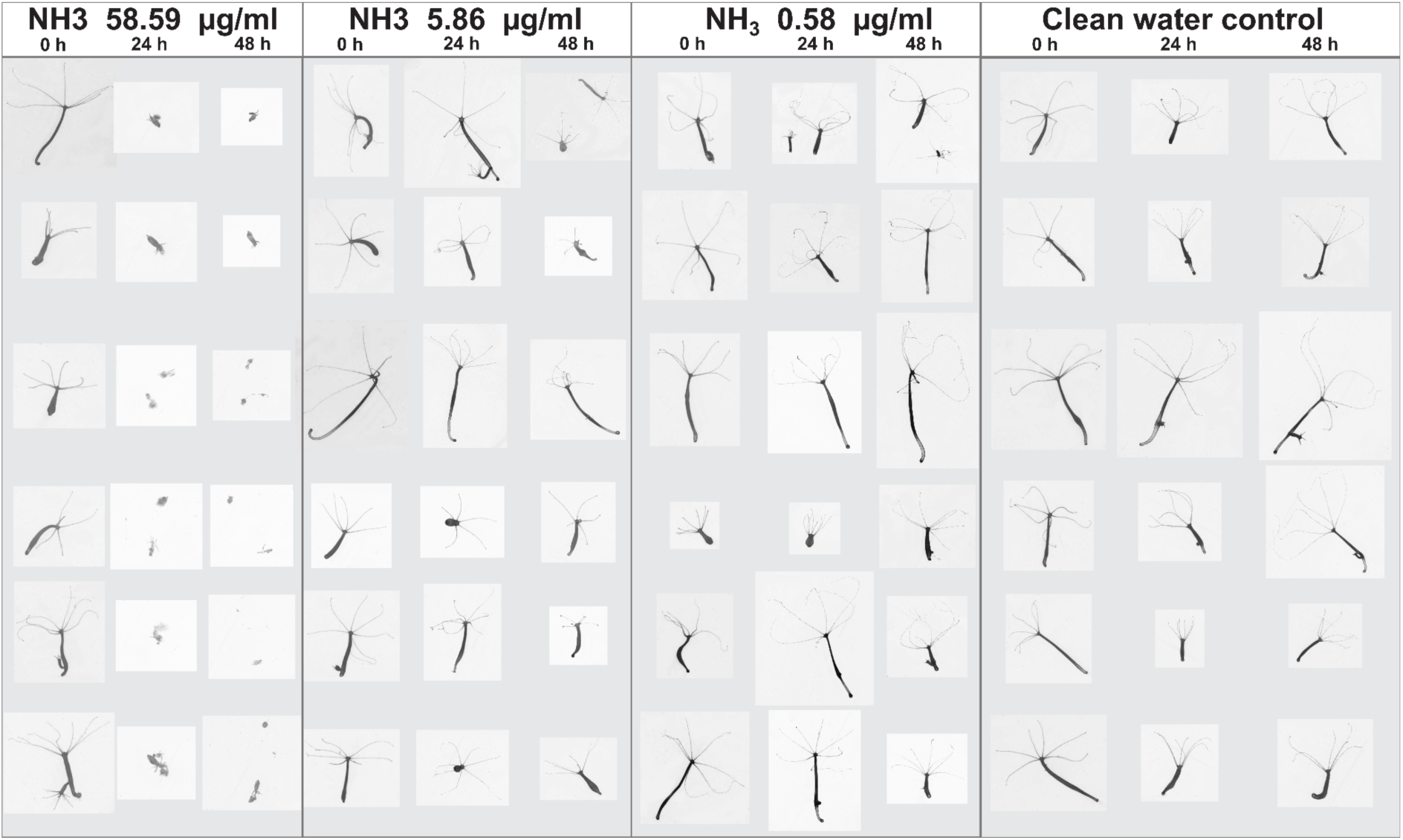
Impact of environmental ammonia on *Hydra* physiology.

