## Supplementary figures and images for "Microbial genome functions explain metabolite-driven dysbiosis and *Pseudomonas*-associated ammonium toxicity in *Hydra*"

### Supplementary File 1

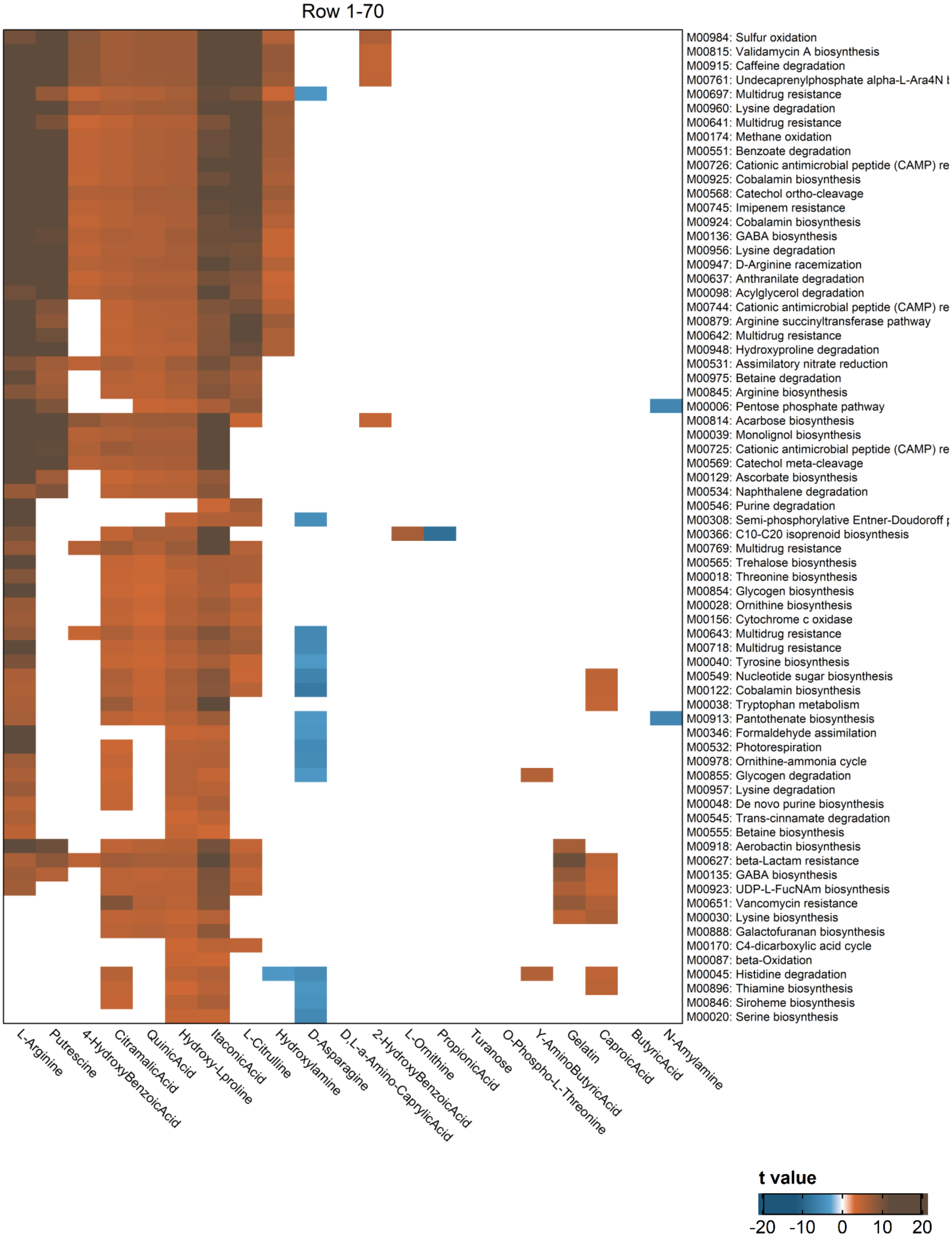

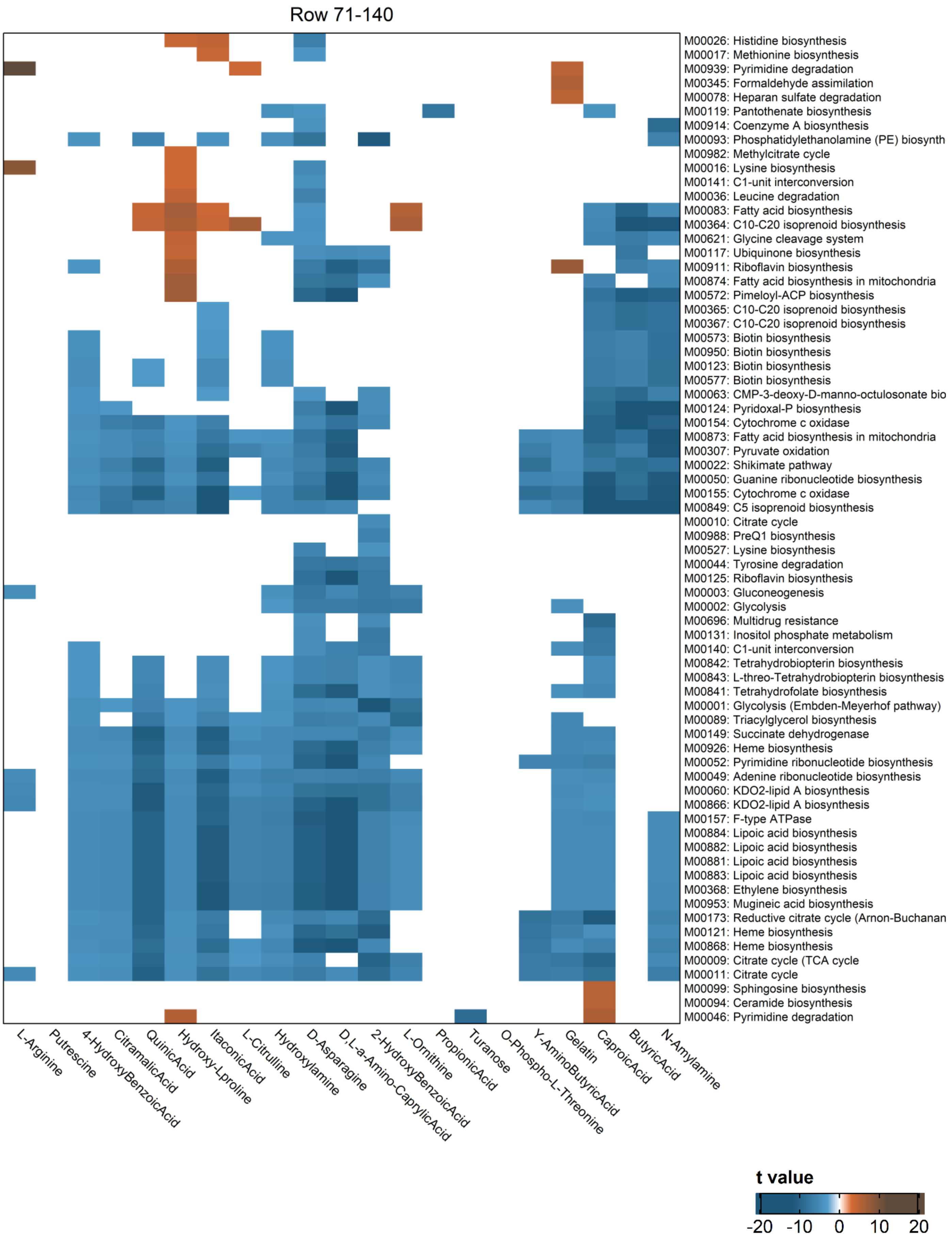

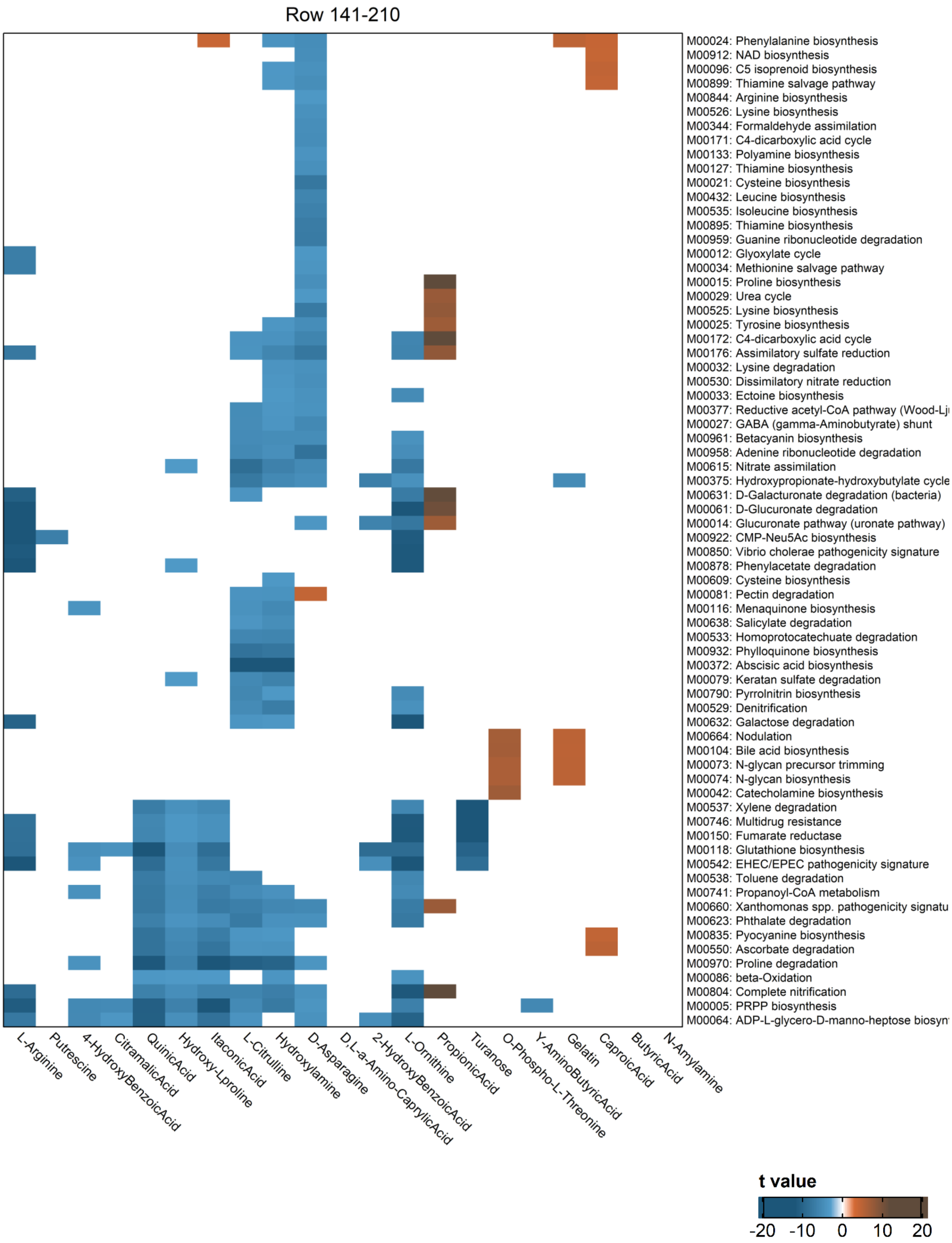

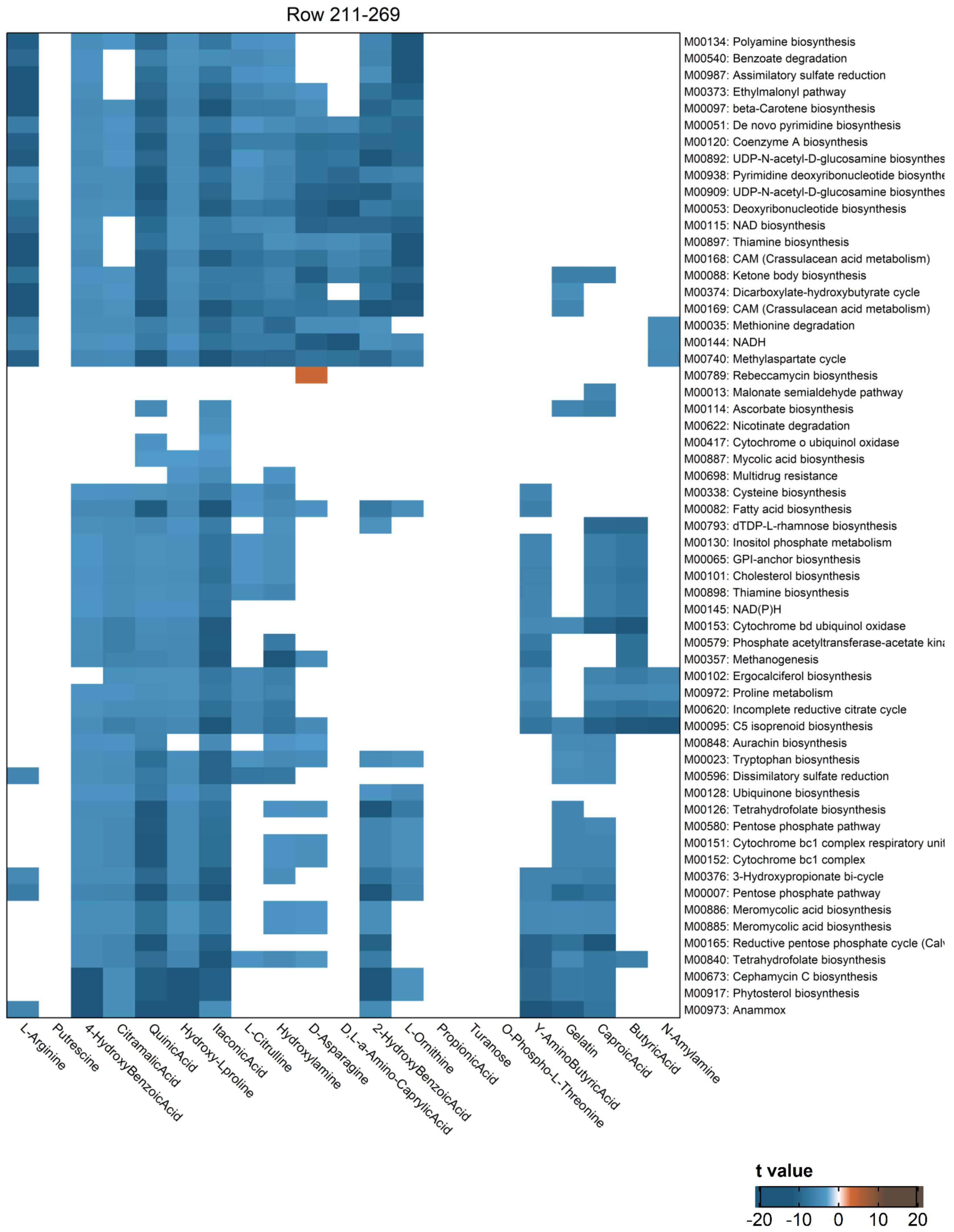
